# PICLN modulates alternative splicing and ensures adaptation to light and temperature changes in plants

**DOI:** 10.1101/2022.06.14.496170

**Authors:** Julieta L. Mateos, Sabrina E Sanchez, Martina Legris, David Esteve-Bruna, Jeanette C. Torchio, Ezequiel Petrillo, Daniela Goretti, Noel Blanco-Touriñán, Danelle K. Seymour, Markus Schmid, Detlef Weigel, David Alabadí, Marcelo J. Yanovsky

**Author notes:** These authors contributed equally (Julieta L. Mateos, Sabrina E Sanchez). Footnote: Dedicated to Dr. Silvia Braslavsky in her 80^th^ anniversary as her amazing journey through science brings her to a memorable landmark.

## Abstract

Plants undergo transcriptome reprogramming to adapt to daily and seasonal fluctuations in light and temperature conditions. While most efforts have focused on the role of master transcription factors, the importance of splicing factors modulating these processes is now emerging. Efficient pre-mRNA splicing depends on proper spliceosome assembly, which in plants and animals requires the methylosome complex. PICLN is part of the methylosome complex in both humans and *Arabidopsis thaliana*, and we show here that the human *PICLN* ortholog rescues phenotypes of *A. thaliana picln* mutants. Altered photomorphogenic and photoperiodic responses in *A. thaliana picln* mutants are associated with changes in pre-mRNA splicing, which partially overlap with those in *prmt5* mutants. Mammalian PICLN also acts in concert with the Survival Motor Neuron (SMN) complex component GEMIN2 to modulate the late steps of UsnRNP assembly, and many alternative splicing events regulated by *PICLN* but not *PROTEIN-ARGININE METHYL TRANSFERASE 5* (*PRMT5*), the main protein of the methylosome, are controlled by *A. thaliana GEMIN2*. As with *GEMIN2* and *SME1/PCP*, low temperature, which increases *PICLN* expression, aggravates morphological and molecular defects of *picln* mutants. Taken together, these results establish a key role for PICLN in the regulation of pre-mRNA splicing and in mediating plant adaptation to daily and seasonal fluctuations in environmental conditions.

## Introduction

Plants sense changes in light and temperature and adjust their growth and development accordingly. They do so, largely, by reprogramming their transcriptome, not only through changes in transcription but also by modulating several co/post-transcriptional processes (Hernando et al., 2017; Dikaya et al., 2021)

An important co/post-transcriptional process that mediates plant adjustment to temperature fluctuations is pre-mRNA splicing, which involves the removal of specific segments (introns) from premature mRNAs, and the ligation of the remaining segments (exons), to produce mature mRNAs. This cut and paste process can take place in alternative ways, leading to skipping or inclusion of specific introns and/or exons, as well as to the extension or shortening of particular segments of pre-mRNA molecules. This process, known as alternative splicing, results in varied mRNA molecules derived from a single gene, which enhances transcriptome and proteome diversity.

Pre-mRNA splicing is catalyzed by the spliceosome, a macromolecular complex comprising five small nuclear ribonucleoproteins (snRNPs, U1, U2, U4–U6) and more than 300 specific proteins (Matera and Wang, 2014). Notably, although UsnRNP assembly can take place autocatalytically *in vitro* with purified Sm proteins and small nuclear RNAs (snRNAs) (Raker et al., 1996), two functional complexes, the Protein Arginine Methyltransferase 5 (PRMT5) and the Survival Motor Neuron (SMN)/GEMIN2 complexes, modulate this process (Meister et al., 2001; Paushkin et al., 2002). Changes in the activity of these snRNP assembly complexes are associated with the onset of cancer or neurological disorders in humans. In mammals, the PRMT5 complex, also known as the methylosome, is composed of PRMT5, PICLN (for Ion Chloride nucleotide-sensitive protein) and WD repeat domain 45 (WD45, also known as methylosome protein 50 [MEP50]). This complex recruits Sm proteins and enhances the transfer of Sm proteins onto the SMN complex via PRMT5 activity, which deposits symmetric dimethylation marks on specific arginines of Sm proteins (Brahms et al., 2001). This last step facilitates the loading of Sm proteins onto snRNAs (Meister et al., 2001; Pellizzoni et al., 2002; Neuenkirchen et al., 2008).

In mammals, in addition to being a methylosome component, PICLN serves as an assembly chaperone that mediates the formation of a ring-shaped particle composed of Sm proteins and PICLN itself. In this particle, RNA binding by Sm proteins is not allowed (Chari et al., 2008) until PICLN is phosphorylated (Schmitz et al., 2021), which allows Sm proteins to be transferred to the SMN complex. This transfer displaces PICLN from the ring particle and the SMN/GEMIN2 complex completes the formation of the snRNPs involved in pre-mRNA splicing.

*PICLN* is conserved across eukaryotes, with orthologs present from yeast (*Saccharomyces cerevisiae*) to humans, including plants such as Arabidopsis (*Arabidopsis thaliana*). However, while mutations in *PICLN* are lethal in mice (*Mus musculus*) and worms (Pu et al., 2000; Furst et al., 2002) plants and yeast cells lacking *PICLN* are fully viable (Barbarossa et al., 2014; Huang et al., 2016). In *Drosophila melanogaster*, PRMT5-mediated methylation of Sm proteins is not essential for snRNP biogenesis, highlighting species-specific differences in this fundamental cellular process (Gonsalvez et al., 2008). Nevertheless, a mutation in *Drosophila arginine methyltransferase 5* (*Dart5*), the fruit fly *PRMT5* ortholog, causes a range of splicing defects (Sanchez et al., 2010). Similarly, a mutation in *PRMT5* in Arabidopsis generates genome-wide alterations in pre-mRNA splicing (Sanchez et al., 2010). In addition, Arabidopsis *prmt5* mutants also show delayed flowering, have altered photomorphogenic responses and have a longer circadian period (Pei et al., 2007; Wang et al., 2007; Hong et al., 2010; Sanchez et al., 2010). Notably, transcripts from the core clock gene *PSEUDO RESPONSE REGULATOR9* (*PRR9*) are aberrantly spliced in *prmt5*, and genetic analysis suggests that the long period phenotype of *prmt5* is at least partially caused by missplicing of *PRR9* (Sanchez et al., 2010). Whether PRMT5 requires PICLN to function in Arabidopsis is still unknown, as well as the molecular function of PICLN in plants.

Here we describe the physiological and molecular roles of *PICLN* during plant development and in response to abiotic stresses. PICLN affects alternative splicing in plants and is important for proper physiological responses to light, temperature, and salt stress. While PICLN interacts with PRMT5 and Sm spliceosomal proteins, it appears to be dispensable for PRMT5-mediated methylation. Still PRMT5 and PICLN have both overlapping and independent functions; while inactivation of *PRMT5* causes more pronounced and widespread perturbations in pre-mRNA splicing than loss of *PICLN, picln* mutants also show a subset of splicing alterations found in *gemin2* but not in *prmt5* mutants. Furthermore, similar to *GEMIN2, PICLN* expression was induced in response to low temperatures and the requirement for PICLN activity increased under cold stress. We propose that PICLN modulates pre-mRNA splicing acting at the interface between the PRMT5 methylosome complex and GEMIN2 to ensure adequate spliceosomal activity under fluctuating or stressful environmental conditions.

## RESULTS

### PICLN plays a role in light-dependent development, flowering time and salt stress responses in Arabidopsis, and its function is partially conserved across kingdoms

To evaluate the role of PICLN in plant growth and development, we characterized two independent T-DNA mutants for *PICLN* (*picln-1* and *picln*-2), harboring a single T-DNA insertion in the third and fifth exon of the gene, respectively (Figure 1A). Neither mutant produced full-length *PICLN* transcripts, suggesting that these mutants are strong alleles (Figure 1B). We first focused our analysis on light responses. Both *picln-1* and *picln-2* mutants are hypersensitive to red light, but not blue light, compared to wild-type seedlings, as evidenced by the stronger inhibition of hypocotyl elongation by red light in the mutants (Figure 1C). Both *picln* mutants flowered later than wild-type plants when grown under an intermediate photoperiod (12-h light/12-h dark cycles), but clearly not later than the wild type under long-day or short-day conditions (Figure 1D, Supplemental Figure S1A). Rather than a late flowering phenotype, one of the two *picln* mutant alleles shows a slight early phenotype under long day conditions (Figure S1). Still, this effect is very mild and only present in one of the two alleles. This phenotype was partially reminiscent of *prmt5* mutants that also flower later than the wild type, although in a photoperiod-independent manner (Pei et al., 2007; Sanchez et al., 2010). However, unlike *prmt5* mutants, and even though *PICLN* transcripts are expressed rhythmically, *picln* mutants exhibited a normal period for rhythmic leaf movements (Supplemental Figure S1B), indicating that their biological functions do not fully overlap. Overall, *picln* mutants show retarded growth and are smaller in size (Figure S1 C-D) as previously reported (Huang et al, 2016). We also tested for functional conservation across kingdoms by heterologously expressing human *PICLN* in the Arabidopsis *picln-1* mutant. Sequence alignment with the human orthologue revealed conservation of specific amino acids and a global homology of ∼50% (Figure 1E). The late flowering phenotype under intermediate photoperiods was rescued in all independent transgenic *picln-1* lines over-expressing human *PICLN* or Arabidopsis *PICLN* (Figure 1F-G). This result supports an inter-kingdom conserved function for PICLN.

**Figure 1.**
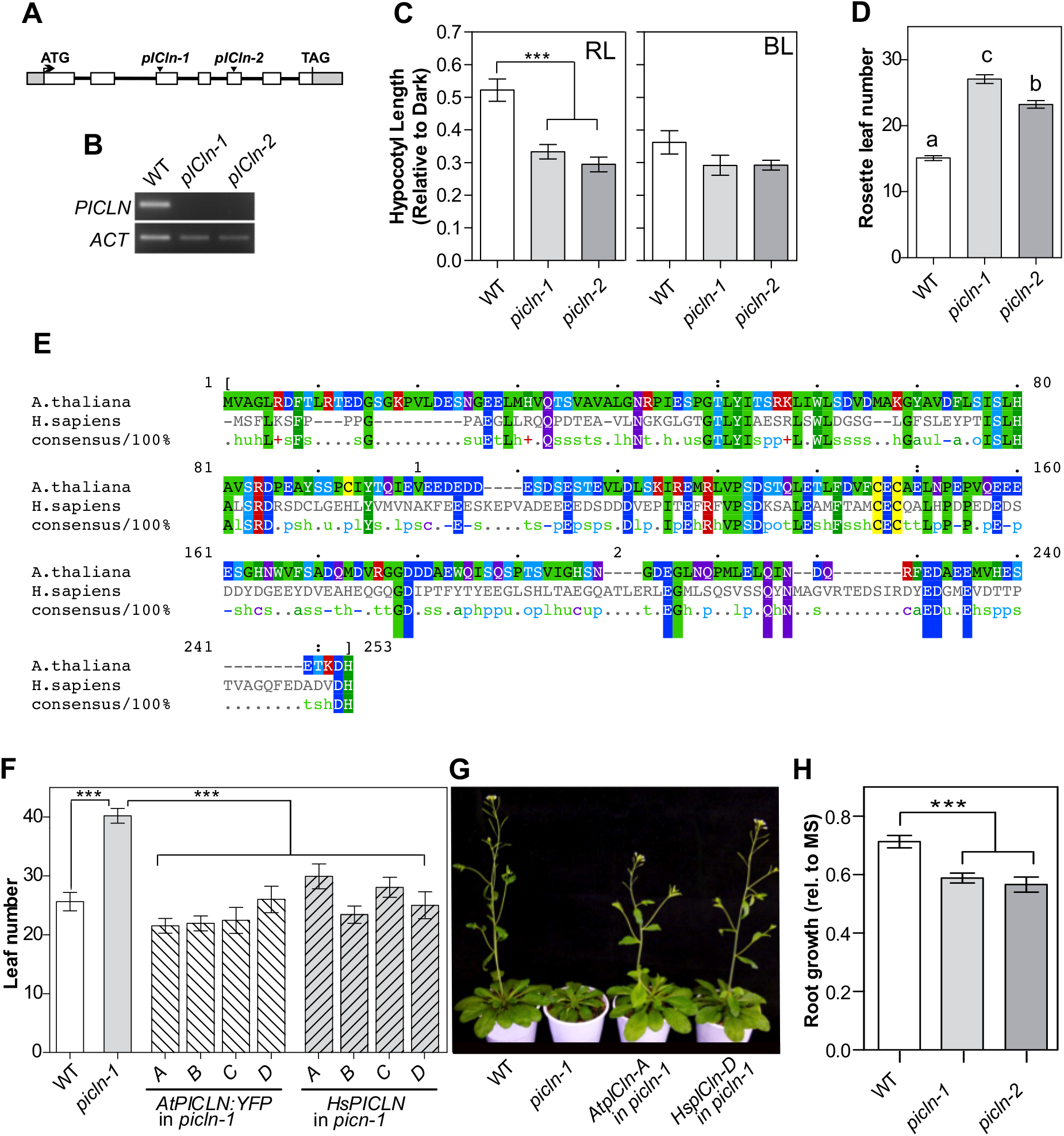
Arabidopsis PICLN shares physiological functions with PRMT5 and its molecular function with human ICln, but mutants are viable. (A) Gene model of Arabidopsis *PICLN* (At5g62290). White boxes, exons; gray boxes, untranslated regions (UTRs); black lines, introns. T-DNA insertion sites in *picln-1* and *picln-2* mutants are indicated with filled triangles. (B) *PICLN* expression in wild-type, *picln-1* and *picln-2* seedlings measured by end-point RT-PCR. *ACT2* was used as a control. (C) Hypocotyl length of seedlings grown under continuous red light (RL) or blue light (BL), normalized to hypocotyl length of etiolated seedlings. (D) Rosette leaf number at flowering time, in plants grown under a 12-h light/12-h dark photoperiod. Different lowercase letters indicate significant difference, as determined by Tukey’s test for multiple comparisons (n>18 plants, *P*<0.0001). (E) Sequence alignment and amino acid conservation of arabidopsis and human PICLN. PICLN sequences in Arabidopsis thaliana (NP_001119478), Homo sapiens (NP_001298128) were obtained from NCBI. (F) Rosette Leaf number at flowering time of wild type, *picln-1*, four independent *picln-1* lines overexpressing either the Arabidopsis (*AtPIClN:YFP)* or human (*HsPICLN) PICLN*. The number of rosette leaves at flowering time was estimated in a 12-h light/12-h dark photoperiod. Significant difference, as determined by Tukey’s test for multiple comparisons (n>16 plants, *P*<0.0001) (G) Representative photograph of the genotypes in (F), illustrating the full rescue of flowering phenotypes of the *picln* mutant by At*PICLN* and Hs*PICLN*. (H) Plant survival in plates containing 160mM NaCl relative to survival in MS media without added salt. Different lowercase letters indicate significant difference, as determined by Tukey’s test for multiple comparisons (n>18 plants, P<0.05)

PICLN was initially thought to be an ion channel that transduces chloride currents during cell swelling (Krapivinsky et al., 1994; Santoro et al., 1998; Gschwentner et al., 1999), although the structural domains and localization of the protein do not support this hypothesis. A subsequent study indicated that PICLN interacts with proteins that respond to hypotonic stress, thereby explaining the observed effects on chloride currents (Dossena et al., 2011). Still, this characteristic does not appear to be conserved in Arabidopsis (Huang et al., 2016). In addition, PRMT5, the core methylosome component, is required for salt stress responses in Arabidopsis (Zhang et al., 2011). Thus, we examined the response of *picln* mutants to salt. Roots of both alleles of *PICLN* showed a greater reduction in growth when grown in a medium containing salt than wild-type plants, indicating that that as with the loss of PRMT5 function, the mutation in *PICLN* confers reduced tolerance to salt stress (Figure 1H).

Together, our findings highlight that PICLN regulates diverse aspects of plant growth and development under varying environmental conditions and this role overlaps only partially with that of its methylosome partner PRMT5.

### *PICLN* is co-expressed and interacts genetically with components of the spliceosome and the spliceosome assembly machinery

As PICLN and PRMT5 are part of the same complex in mammals and yeast, we reasoned that their encoding transcripts might be co-expressed in plants at different developmental stages to support their shared functions. Examination of publicly available datasets indicated that *PICLN* and *PRMT5* are co-expressed across many conditions in Arabidopsis (Figure 2A). *PICLN* expression was also positively correlated with that of *SME1* (also known as *PORCUPINE* [*PCP*]) (Capovilla et al., 2018; Huertas et al., 2019) and *GEMIN2* (Schlaen et al., 2015) (Figure 2B-C). This co-regulation between *PICLN* and genes encoding components of the spliceosome assembly machinery and the spliceosome itself across a wide range of conditions suggested that the encoded proteins might function non-redundantly in the same molecular process(es). We explored this possibility by characterizing the genetic interactions between *PRMT5, SME1/PCP* and *GEMIN2* with *PICLN*, in double mutant plants. Genetic analyses revealed severe synergistic defects in the *picln-1 prmt5-5* double mutant: embryos were arrested at the late globular stage showing in some cases more than one layer of cells within the suspensor (Figure 2D). Similarly, we failed to identify any double mutant among the F_2_ progeny of a cross between *picln-1* and *gemin2-1* (Schlaen et al., 2015) or *picln-1* and *sme1-1* (Supplemental Figure S3), suggesting that the simultaneous loss of these proteins is embryo-lethal (Figure 2E, F). Analysis of embryonic phenotypes in seeds taken from the siliques of plants harboring a single functional copy out of four indicated that again both *picln-1 sme1-1* and *picln-1 gemin2-1* double mutant embryos were arrested at the late globular stage and show aberrations in the suspensor. We obtained similar synergistic genetic results between *picln* and *smd1a* or *smd3b* mutants (Supplemental Figure S3). These results support the idea that PICLN could act in concert with PRMT5 and GEMIN2 to modulate snRNP assembly in Arabidopsis.

**Figure 2.**
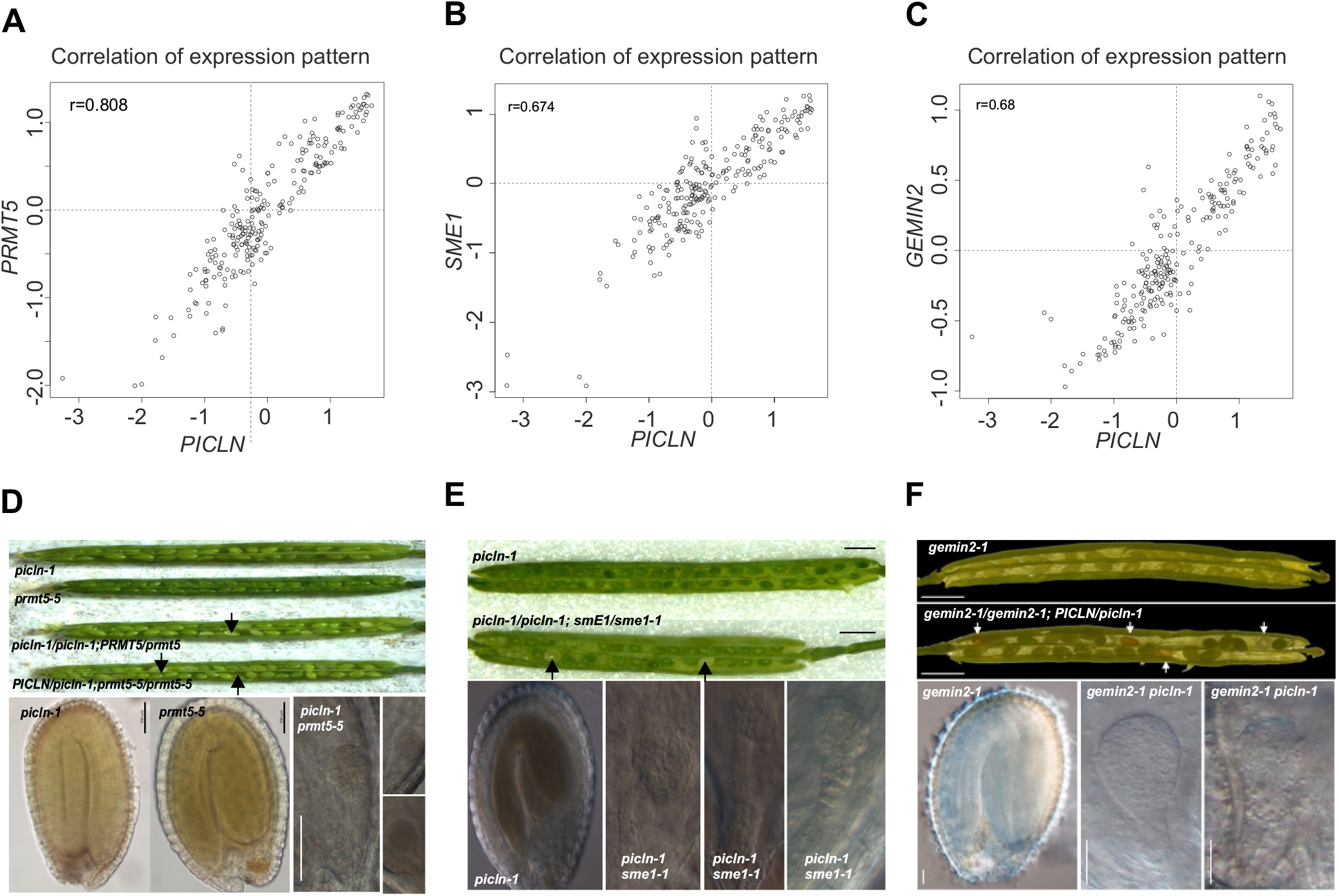
*PICLN* is co-expressed and interacts genetically with components of the spliceosome assembly machinery. (A-C) Correlation of expression patterns between *PICLN* and *PRMT5* (A), *SmE1/PCP* (B) and *GEMIN2* (C). Both axes are relative gene expression values (Log_2_-normalized) against the averaged expression levels of each gene. Data obtained from http://atted.jp/. (D-E) Embryonic phenotypes of single and putative double mutants for *picln-1* and *prmt5-5* (D), *sme1-1/pcp-1*(E) and *gemin2-1* (F). Pictures were taken from siliques dissected 45 to 60 days after sowing. Top: Dissected siliques showing developing seeds for the indicated genotypes. Arrows indicate some of the aborted putative double mutant seeds. Bottom: DIC images of seeds from homozygous or heterozygous genotypes.

### The *picln* mutant has global defects in alternative splicing

The high correlation in gene expression with genes encoding spliceosome assembly components known to modulate alternative splicing prompted us to investigate the participation of PICLN in pre-mRNA splicing. To this end, we performed RNA-seq experiments to compare the transcriptome of the *picln-1* mutant with that of wild-type seedlings. We evaluated three biological replicates from seedlings grown for 12 days under 12-h light/12-h dark cycles, a condition in which we observed the late flowering phenotype in the mutant. We focused on the *picln-1* allele, as both alleles behaved similarly (Figure 1, Supplemental Figure S1). We identified 278 differentially expressed genes (DEGs) between *picln-1* and wild-type seedlings (Supplemental Table 1). When we looked at the expression of other components of the methylosome and splicing machinery such as the *SM* genes, we found no difference in expression between genotypes, except for a mild change in the levels of *SME1/PCP* (Supplemental Figure S2). mRNA levels of *GEMIN2* and *PRMT5* also accumulated similarly in *picln-1* and wild-type plants. Recently, Willems and co-workers showed that the levels of the U2snRNA but not of U1, U4 and U5 snRNAs, are altered in *picln* mutants compared to wild-type plants (Willems et al., 2022). Among DEG genes, we noticed genes that contribute to flowering time regulation, namely *CYCLING DOF FACTOR1* (*CDF1*) (Fornara et al., 2009), *SQUAMOSA LIKE-PROTEIN3* (*SPL3*) (Yamaguchi et al., 2009), and *GA REQUIRING 2/ENT-KAURENE SYNTHASE* (*GA2/KS*) (Yamaguchi et al., 1998). In addition, we observed downregulation of *HYPOCOTYL IN FAR-RED* (*HFR1*), a gene involved in photomorphogenic responses, in *picln* mutants. Whether any of these alterations in transcript levels are responsible for the flowering or hypersensitivity to red light phenotypes of the *picln-1* mutant needs further research. We also conducted a gene ontology (GO) analysis of the DEGs, which revealed an enrichment for genes related to the regulation of defense responses (Supplemental Figure S2). This result was in agreement with the previous observation that oomycete spores have a reduced growth in *picln* mutants (Huang et al., 2016).

We then analyzed the effects of *picln-1* on pre-mRNA splicing. To this end, we evaluated four different types of splicing events: alternative usage of 5′and 3′splicing sites (alt 5′; alt 3′), intron retention (IR); and exon skipping (ES) using the ASpli package (Mancini et al., 2021). We then compared the relative distribution of the different types of AS events among all annotated AS events in the genome, with the relative distribution of the different types of AS events among all the events that were differentially spliced in *picln* mutants compared to wild-type plants (Figure 3A). For this, we analyzed and scored all possible splicing events from 20,329 genes expressed above a minimal threshold level in both wild-type and *picln-1* seedlings. We detected the differential usage of 541 splicing sites in the *picln-1* mutant relative to the wild type (Supplemental Table S1). Among splicing events affected in the *picln-1* mutant, we observed IR events as the most affected (Figure 3A). We independently validated two events belonging to different AS classes by RT-PCR (Supplemental Figure S5). GO analysis using all differentially spliced genes showed that most are related to RNA metabolism (Figure 3B). Interestingly, genes related to mRNA polyadenylation were strongly overrepresented in the *picln-1* mutant (over twenty-fold relative to their frequency in the Arabidopsis genome), highlighting a putative role for *PICLN* in controlling splicing of genes that regulate gene expression and RNA processing. Strikingly, some genes involved in the circadian clock also exhibited defects in their splicing patterns in the mutant. We detected retention of intron 4 of *TIMING OF CAB2 EXPRESSION 1* (*TOC1*), and of intron 7 of *PSEUDO RESPONSE REGULATOR 3* (*PPR3*). Despite these alterations in two genes encoding core clock components, we did not observe any circadian phenotype in the *picln-1* mutant (Supplemental Figure S1). We also observed significant retention for intron 2 of the *PHOTOTROPIN 2* (*PHOT2*) transcript and for intron 2 of *FAR-RED ELONGATED HYPOCOTYLS* (*FHY3*) (Supplemental Table S1), which could be contributing to the altered photomorphogenic behavior observed in *picln-1* mutants (Figure 1C.) We then evaluated the frequencies of nucleotide sequences around the 5′ splice sites and the 3′splice sites for all IR events affected in *picln-1* compared to the consensus 5′ splice site or 3′splice site of all introns. We found no variation at the donor or acceptor splice site sequence (Figure 3C), in contrast to our previous results with the *prmt5* mutant, in which the consensus A and G nucleotides at positions −2 and – 1 of the donor site were underrepresented (Sanchez et al., 2010). Together, these results show that PICLN is involved in regulating splicing and suggest that the underlying mechanism is at least partially distinct from that of PRMT5.

**Figure 3.**
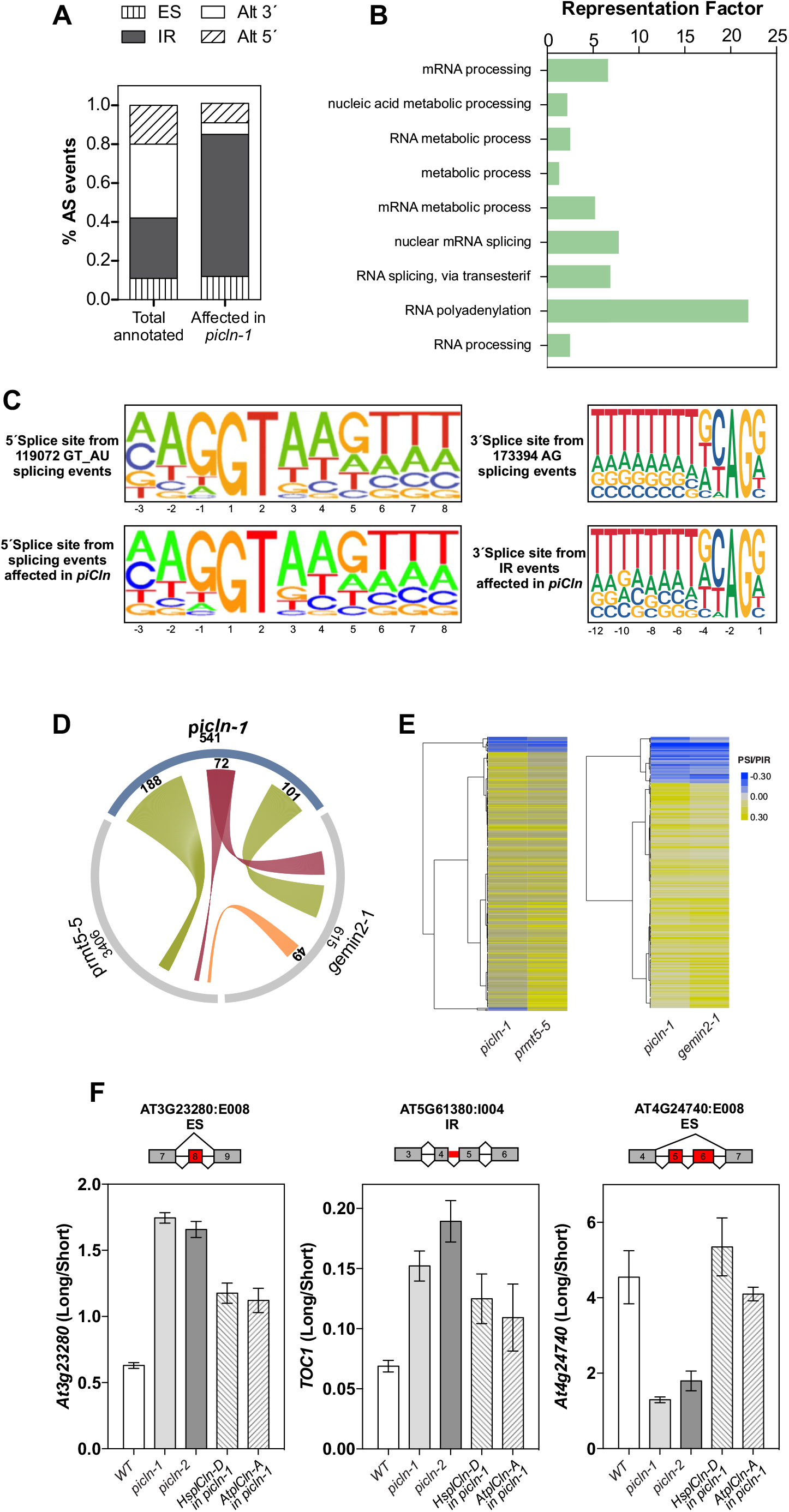
The *picln* mutant shows global defects in alternative splicing, some shared with PRMT5, SmE1/PCP and GEMIN2, and can be rescued with Human as well as Arabidopsis PICLN. (A) Relative frequencies of different alternative splicing (AS) types for all detected AS events; Alt 3’ and Alt 5’, alternative acceptor and donor splice sites, respectively; ES, exon skipping, IR, intron retention. (B) Gene Ontology-term enrichment analysis of genes whose splicing is affected in *picln-1*. (C) (Left) Pictogram logos showing the frequency distribution of nucleotides at the 5′ splicing site of 119,072 GT_AG_U2 Arabidopsis introns or of the 541 splicing events significantly altered in *PICLN*. (Right) Pictogram logos showing the frequency distribution of nucleotides at the 3′ splicing site of 173,394 AG Arabidopsis introns or of the 541 splicing events significantly altered in *PICLN*. (D, E) Comparison of the splicing defects detected in *picln-1, gemin2-1* and *prmt5-5* single mutants. (D) Summary of the analysis, shown as a Circos plot. The total number of affected events in each mutant is shown below the mutant name. Connecting lines are scaled and represent shared affected events. Red lines, shared across all genotypes; green lines, shared between *picln-1* and one of the other mutants; orange lines, shared between *gemin2-1* and *prmt5-5*. Numbers above the lines represent the number of events. (E) Hierarchical clustering of 260 common events between *picln-1* and *prmt5-5* (left) or 173 shared between *picn-1* and *gemin2-1* (right). Clustering was done with the complete linkage method with Clustal 3.0 Software. PSI or PIR (percent intron retention/percent spliced-in) values were calculated to measure the extent of the splicing defect. (F) Example of a splicing event affected in two *picln* alleles and complemented either with *HsPICLN* or *AtPICLN*. Validation was done by RT-PCR quantifying the long and short isoform and showing the ratio. Gene scheme with exons as boxed and introns as lines, the

### Splicing alterations in *picln* mutants are partially shared with *prmt5* and *gemin2* mutants

The snRNPs mainly function as core components of the spliceosome. Indeed, seven Sm proteins associate to form an heptameric ring that contains a snRNA (U1, U2, U4, U5). During this assembly, Sm proteins that are methylated by PRMT5 in mammals (Friesen et al., 2001a) are assisted by PICLN, which acts as a chaperone to prevent binding to non-specific RNAs at this early stage of snRNP assembly. At a later stage, a transfer of PICLN-bound Sm proteins to the SMN complex provokes the displacement of PICLN from the complex, thereby allowing snRNA binding, ring closure, and snRNP release. To better understand how PICLN and PRMT5 affect splicing in plants, and to evaluate if their molecular functions overlap, we performed an RNA-seq in the *prmt5-5* mutant and compared its effect on gene expression and splicing with that of *picln-1*. Overall, we identified 1,500 genes with altered expression in *prmt5-5* compared to wild-type seedlings, while the expression of only 278 genes was affected in the *picln-1* mutant, 122 of which were also altered in *prmt5-5* (Supplemental Figure S6, Supplemental Table S1). This observation suggests that PRMT5 has a larger role in regulating gene expression than PICLN. The *picln-1* mutation caused ∼60% of the DEGs to be downregulated relative to wild-type seedlings, and compared to ∼45% of DEGs in *prmt5-5*. More importantly, for 117 out of 122 DEGs whose expression was altered in both *prmt5-5* and *picln-1* mutants, the direction of misexpression was the same, underscoring that both components generate the same effect on the transcriptome (Supplemental Figure S6). Characterizing the contributions of PICLN and PRMT5 to alternative and constitutive splicing again revealed a broader effect of *prmt5-5* compared to that of *picln-1*. As seen for transcript abundance, half of the splicing events affected in *picln-1* (541 in total) were also affected in *prmt5-5* (Supplemental Figure S6). In addition, we identified examples of splicing defects specific to each genotype (Supplemental Figure S7). We validated several of these altered events, both shared and genotype-specific splicing defects, by end-point RT-PCR (Supplemental Figure S8). Analyzing the effect of each mutation separately, we determined that only 4% of the DEGs in *picln-1* were affected in splicing, while approximately 25% of the DEGs in *prmt5* also had a defect in splicing. This result suggests that splicing and transcription alterations are at least partially coupled for PRMT5, but this effect is less relevant for PICLN.

To further characterize similarities and differences between the effects on splicing mediated by PICLN and those of other proteins involved in snRNP assembly, we also compared splicing events affected in *picln-1* and *prmt5-5* with those found to be altered in the *gemin2-1* mutant in a previous work (Schlaen et al., 2015). A genome-wide evaluation indicated that 30% of the splicing defects observed in *picln-1* are shared with those found in *gemin2-1* while, as mentioned above, approximately 50% of the splicing events altered in *picln-1* were also affected in *prmt5-5* (Supplemental Figure S5, Figure 3D). Interestingly, only 12% of the events affected in *picln-1* (72 splicing events involving 64 genes) were simultaneously affected in both the *prmt5-5* and *gemin2-1* mutants (Figure 3D, red links). To note is that neither mRNA levels of *PRMT5* nor *GEMIN2* were changed in the absence of *PICLN* (Supplemental Figure S2). Besides, the mutation in *PICLN* did not affect the splicing of these two genes (Supplemental Table S1). All these results suggest that Arabidopsis PICLN participates in splicing in two different ways: as an adaptor protein of the PRMT5 methylosome complex and as assembly chaperone of the GEMIN2/SMN complex.

A particularly interesting example of a difference between PICLN and PRMT5 effects on splicing was that of *U1-70K*. This gene encodes a key splicing factor whose mRNA is alternatively spliced in Arabidopsis and generates one of two forms: a functional isoform that encodes the full-length protein, and an alternatively spliced isoform that retains intron 6 and results in a truncated protein that interferes with U1 snRNP functionality (Golovkin and Reddy, 1996). Strikingly, we detected increased amounts of the fully spliced functional U1-70K isoform in the *picln-1* and *gemin2-1* mutants compared to wild-type seedlings (Supplemental Figure S7D), likely reflecting a homeostatic mechanism that compensates for the attenuated activity of the spliceosomal machinery in these mutants. By contrast, we observed substantial retention of intron 6 in the *prmt5-5* mutant, indicating that PICLN and GEMIN2 share a common effect on splicing, which is at least partially different from that of PRMT5 (Supplemental Figure S7D).

The effect of a mutation on splicing can be quantified by comparing PSI (percent of inclusion) and PIR (percent of IR) values of AS events in different genotypes (wild-type, *pciln-1, prmt5-5* and *gemin2-1*). Hierarchical clustering of PSI/PIR values associated with splicing alterations shared between *picln-1* and *prmt5-5*, and between *picln-1* and *gemin2-1*, showed a similar trend in the mutants (Figure 3E, Supplemental Figure S7). This indicates that for common altered AS events PICLN has a similar effect as PRMT5 or GEMIN2.

Encouraged by the observation that introducing human *PICLN* into Arabidopsis can rescue the late flowering phenotype of *picln-1* (Figure 1F-G), we wished to test whether human *PICLN* would similarly restore the splicing defects observed in Arabidopsis *picln* mutants. Indeed, human *PICLN* rescued the splicing defects seen in the Arabidopsis *picln-1* and *picln-2* mutants (Figure 3F) to a similar extent as Arabidopsis *PICLN* does, further supporting the conservation of PICLN molecular function across kingdoms.

### PICLN interacts with several Sm proteins but Sm symmetric dimethylation is not affected in *picln* mutants

In mammals, the methylosome complex comprises PRMT5, PICLN, and WD45 (Friesen et al., 2001b; Chari et al., 2008). PICLN was suggested to confer specificity to PRMT5 for methylating Sm proteins, which facilitates spliceosome assembly (Pesiridis et al., 2009). Here we tested for direct interactions between PICLN and a series of Sm proteins. In a yeast two-hybrid assay (Y2H), we detected a strong interaction between PICLN and each of SmBb, SmD1 and SmD3b (Figure 4A), weaker interactions with SmE1/PCP and SmF, and no interaction with SmG (Figure 4A). In humans, Sm proteins contain multiple arginine-glycine (RG) repeats in their C termini that are methylated by PRMT5 (Branscombe et al., 2001; Friesen et al., 2001b; Meister et al., 2001). We therefore analyzed the sequences of Arabidopsis Sm proteins and found that as their human orthologs, they also contained RG repeats, which could be substrates for symmetric dimethylation (Supplemental Figure S8). To test if the Sm proteins that interact with PICLN were symmetrically methylated at their R residues, we generated transgenic lines overexpressing constructs encoding SmBb, SmD1 or SmD3 fused to different fluorescent proteins (SmB-YFP [yellow fluorescent protein], SmD1-CFP [cyan fluorescent protein], and SmD3-GFP). As a positive control, we used plants overexpressing a tagged version of SM-LIKE PROTEIN 4 (LSM4), a protein previously shown to be symmetrically dimethylated by PRMT5 (Deng et al., 2010).

**Figure 4.**
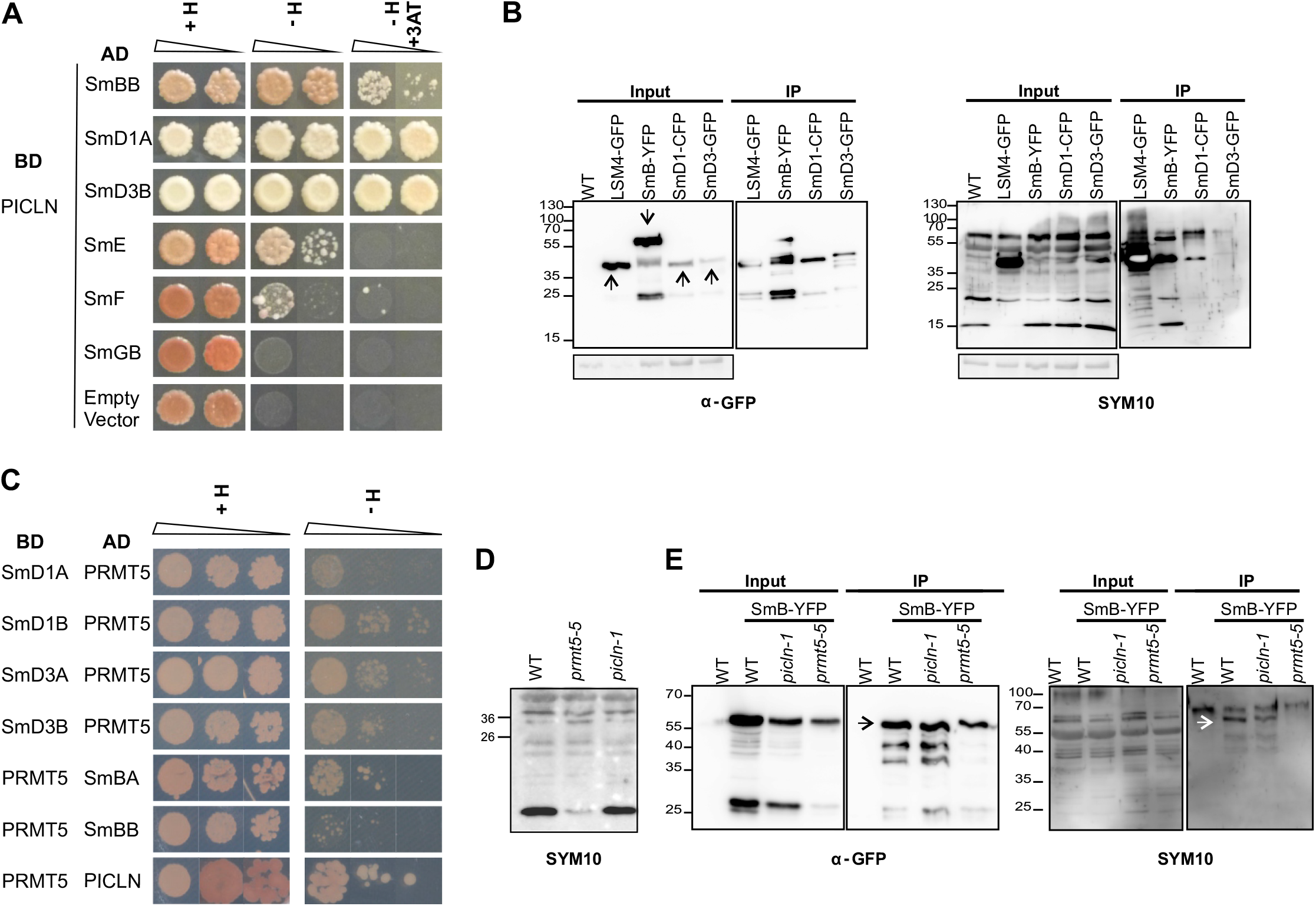
PICLN interacts with several Sm proteins but Sm symmetrical dimethylation is not affected in *PICLN*. (A) Yeast two-hybrid assay of the interactions between PICLN and the different subunits of the Arabidopsis Sm complex. Two serial dilutions per combination of constructs are shown. +H, control medium containing histidine; –H and –H+3AT, synthetic defined (SD) medium lacking histidine and also containing 5 mM 3-amino triazole (3-AT), respectively. Pictures were taken after 6 days of growth at 28ºC. (B) Sm proteins that interact with PICLN are symmetrically dimethylated. Immunoprecipitation (IP) assays in fluorescent protein-tagged *Sm* overexpressing lines. (SmB-YFP, SmD1-CFP, SmD3-GFP and LSM4-YFP with anti-GFP antibody. Immunoblots with anti-GFP and anti-dimethyl arginine antibody, symmetric, SYM10 antibody. LSM4-YFP plants were used as a positive control. Ponceau red staining shows equal loading across samples. Arrows represent expected size. (C) PRMT5 can interact with Sm proteins and PICLN in yeast. Y2H assays of interactons between PRMT5 and Sm or PICLN proteins. Pictures are from yeasts grown on -WL and -WLH selective media for additional 4 days before taking pictures. (D) Methylation of Sm proteins is not affected in *picln* mutants. Immunoblot of wild-type, *prmt5-5* and *picln-1* samples with the SYM10 antibody. The *prmt5-5* mutant was used as a positive control for impaired methylation. (E) SmB methylation is not altered in *picln-1*. IP of SmB-YFP in wild-type, *picln-1* and *prmt5-5* background using anti-GFP antibody. Western blots were performed with anti-GFP and SYM10 antibodies. Arrows show the expected size for SmB-YFP.

We then performed immunoprecipitation assays using an anti-GFP antibody, and evaluated symmetric dimethylation with the anti-dimethyl-arginine antibody, symmetric (SYM10). In all cases, we observed the symmetric dimethylation of Sm proteins (Figure 4B). We further explored the interaction of Sm proteins with PRMT5, the other component of the methylosome by Y2H. We found that SmD1A, SmD1B, SmD3A, SmD3B, SmBA and SmBB specifically interact with PRMT5 (Figure 4C, Supplemental Figure S10). Moreover, we validated the previous interaction between PRMT5 and PICLN (Figure 4C). Remarkably, global symmetric arginine dimethylation was not affected in *picln-1* mutants (Figure 4D) while, as expected, *prmt5-5* extracts showed reduced methylation (Figure 4D). In a similar manner, symmetric dimethylation of SmB:YFP in *picln-1* was almost unchanged (Figure 4E). Accordingly, subcellular localizations of SmB, SmD1 or SmD3 were not changed in the absence of *PICLN* (Supplemental Figure S11). Sm proteins can still translocate to the nuclei and are probably capable of forming snRNPs. On the other hand, *PICLN* can localize in the nucleus as well as in the cytoplasm (Supplemental Figure S11).

### PICLN regulates low temperature responses and growth

To adjust to changing environmental conditions, plants dynamically regulate the transcriptional output of genes related to stress responses. However, changes in the environment also influence post-transcriptional regulation of mRNAs, such as pre-mRNA splicing, mRNA degradation and stability (Floris et al., 2009; Nakaminami and Seki, 2018) to shape the transcriptome and allow plant adaptation. Recently, several splicing-related factors, including GEMIN2 and SME1/PCP, have been shown to be regulated by temperature and to contribute to the adaptive response of plants to temperature variations (Schlaen et al., 2015; Okamoto et al., 2016; Carrasco-López et al., 2017; Capovilla et al., 2018; Catalá et al., 2019; Huertas et al., 2019). We therefore asked whether PICLN was also involved in modulating temperature responses. When kept at 10°C under continuous light for 8 weeks, we observed that the *picln-1* mutant, but not wild-type plants, showed a strong reduction in overall growth (Supplemental Figure S12). Furthermore, we noticed a more pronounced inhibition of root elongation at 10°C compared to warm conditions in *picln-1* seedlings relative to the wild-type (Figure 5A-B). *PICLN* transcripts rose two-fold after exposing seedlings to 10ºC for 24 h (Figure 5C). Most snRNP components did not show a significant change in expression levels upon cold treatment, with the exception of *SME1* and SMD3A, which changed 2.7 and 2.0 fold respectively (Supplemental Figure S12). These results suggest that PICLN is part of a regulatory mechanism that contributes to adjusting plant growth to changes in temperature conditions.

**Figure 5.**
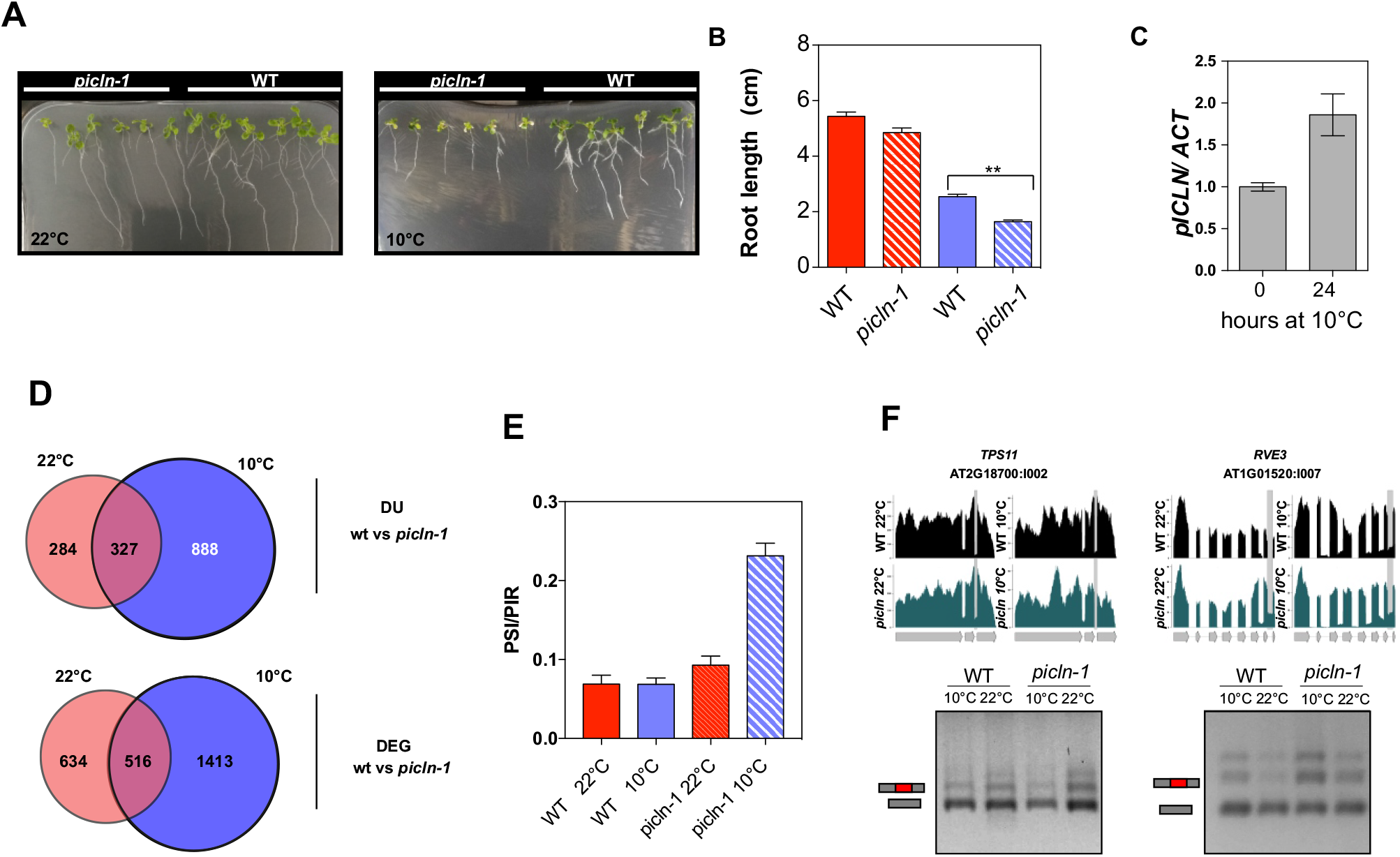
*PICLN* modulates low temperature effects on alternative splicing and growth in Arabidopsis. (A) Cold sensitivity of wild type and *picln-1* at 10°C. Four-day-old seedlings grown at 22°C in a 16-h light/8-h dark photoperiod were kept at 22°C as control or transferred to continuous light at 10°C. Pictures were taken 3 weeks or 8 weeks later, respectively. (B) Root growth on MS medium at 22°C (red bars) or 10°C (blue bars). Primary root length of wild type and *picln* mutants was determined 20 d after seedlings were transferred to cold (n=12, *** *P* <0.05) or after 10 d when grown at 22°C. (C) Accumulation of *PICLN* transcripts in 10-d-old wild-type seedlings grown under control conditions at 22°C or exposed to 10°C for 24 h. (D) Venn diagrams showing the extent of overlap in splicing defects (top) and differentially expressed genes (bottom) identified in *picln-1* in comparison to wild-type plants at 22°C or after a shift to 10°C for 24 h. (E) Average change in PSI/PIR values for all genes for pre-mRNA splicing events that respond to temperature either in wild-type or the *picln-1* mutant. Error bars represent SEM. (F) Integrated genome browser view (top) of coverage plots from two genes with splicing defects that resulted from the interaction between genotype and temperature. Bottom: validation of splicing defects via RT-PCR.

To investigate the molecular consequences of the loss of PICLN function at this temperature, we conducted an RNA-seq analysis on wild-type and *picln-1* mutant seedlings grown for 10 d at 22°C and then exposed or not at 10°C for 1 d. We discovered that at the lower temperature, PICLN controls twice as many splicing events as at ambient temperature (Figure 5D). Indeed, 554 genes displayed 611 splicing defects in the *picln-1* mutant at 22°C, while this number increased to 1,020 genes with 1,215 splicing alterations in seedlings subjected to 10°C for 24 h (Supplemental Table S1, Figure 5D). Similarly, we detected 1,929 DEGs at 10°C in the mutant, compared to 1,150 genes at 22°C. This analysis clearly revealed that PICLN contributes more strongly to transcriptional and post-transcriptional gene regulation at 10°C than at 22°C. In agreement with this idea, we found that events that were strongly responsive to low temperatures in *picln-1* were almost not responsive in wild-type seedlings (Figure 5E). We also detected differences in splicing events for 65 genes, which were associated with interactions between genotype and temperature conditions, as some AS events altered in the *picln-1* mutant at low temperatures were not affected in the mutant at 22°C (Figure 5F). This result is in line with our hypothesis that PICLN is particularly important to maintain proper splicing of transcripts in seedlings exposed to stressful low temperature conditions.

To gain further insight into the mechanism by which PICLN influences adaptive responses to colder temperatures, we further analyzed those genes with splicing events modulated by PICLN in response to lower temperature conditions. Around 38% of the genes whose AS patterns were affectted by PICLN under cold conditions displayed AS alterations in response to cold treatment in wild-type seedlings (RF:1.9, *P* <1.43 × 10^−21^). In addition, ∼47% of genes whose splicing was regulated by PICLN at 10°C showed changes in their expression when wild-type seedlings were shifted from 23°C to 16°C for 3 d (Capovilla et al., 2018). Interestingly, 172 genes showed simultaneous alterations in splicing and expression in *picln-1* compared to wild-type seedlings at 10°C (RF:3.2, *P* < 5.9 × 10^−44^). Finally, we identified 523 genes with altered expression in *picln-1* mutant seedlings exposed to cold conditions that also responded to the low temperature treatment in wild-type seedlings. Taken together, we conclude that PICLN exerts an important role in modulating gene expression and splicing patterns in response to low temperature in Arabidopsis.

### PICLN is required to control the alternative splicing of splicing factors that respond to salt

PICln was first described as a nucleotide-sensitive chloride conductance regulatory protein. In humans, pICln can form ion channels (Furst et al., 2002) though in Arabidopsis this characteristic seems not to be conserved (Huang et al., 2016). Still, Huang et al., detected pICln in the ER membrane suggesting that it could form an ion channel in this organelle. Besides, PRMT5 is required for proper salt stress responses in Arabidopsis (Zhang et al., 2011) and we observed that *picln* mutants were more sensitive to salt concentrations (Figure 1 H). Therefore, we decided to examine salt related phenotypes in *picln* mutants in more detail, as well as possible molecular mechanisms associated with these phenotypes. The growth of *picln-1* and *picln-2* mutant plants was completely inhibited by 120 or 160 mM NaCl, a behavior that is similar to that of *prmt5-5* mutants (Figure 6A-B). By contrast, wild-type plants showed a higher survival rate. Interestingly, salt exposure did not cause any change in *PICLN* expression (Figure 6C). Nevertheless, salt treatments were enough to promote expression of *RESPONSIVE TO DESICCATION 29A* (*RD29A*), a gene known to respond to cold and salt stress (Figure 6C). It was previously shown that several AS events in salt responsive genes are promoted by salt treatment (Feng et al., 2015). We then contrasted transcriptomic data of wild-type seedlings under salt treatment available from Feng et al (Feng et al., 2015) with that of *picln-1* mutants analyzed here. We found that ∼46% of misregulated genes in *picln-1* were salt-responsive genes (Figure 6D). Most interestingly, 132 genes that were changed upon salt exposure showed splicing defects in *picln-1*, suggesting that *picln-1* salt hypersensitivity might be due to misregulation and differential usage of genes that respond to salt stress (Figure 6D). Interestingly, we found that *PICLN* regulates splicing of a subset of SR proteins and splicing related factors that are also affected by salt stress (Figure 6E). It has been reported that *SR30* generates at least five different transcripts, *SR30* mRNA1–5 (Palusa et al., 2007). At the same time, Tanabe and co-workers showed that salinity at 250 mM NaCl for 1 d resulted in a decrease in the levels of *SR30* mRNA4 (Tanabe et al., 2006). We validated this decrease in the levels of mRNA4 isoform of *SR30* using a salt treatment similar to that performed in Feng et al. (Figure 6E). The alternative 3′ss event was decreased in *picln-1* as well as in salt treatment. In summary, *picln-1* phenotype under salt might be the results of AS defects in salt responsive genes provoked by lack of *PICLN* and particularly of mispliced versions of splicing factors such as SR proteins found in *picln-1*. To some extent, for a subset of genes, *picln-1* mimics the effect of salt on the transcriptome (Figure 6E). Taken together, these results indicate that PICLN is important for proper adaptive responses to multiple abiotic stresses. However, the observation that cold but not salt stresses modulate *PICLN* expression, suggests that *PICLN* is more directly involved in the adaptive responses to a specific subset of abiotic stress.

**Figure 6.**
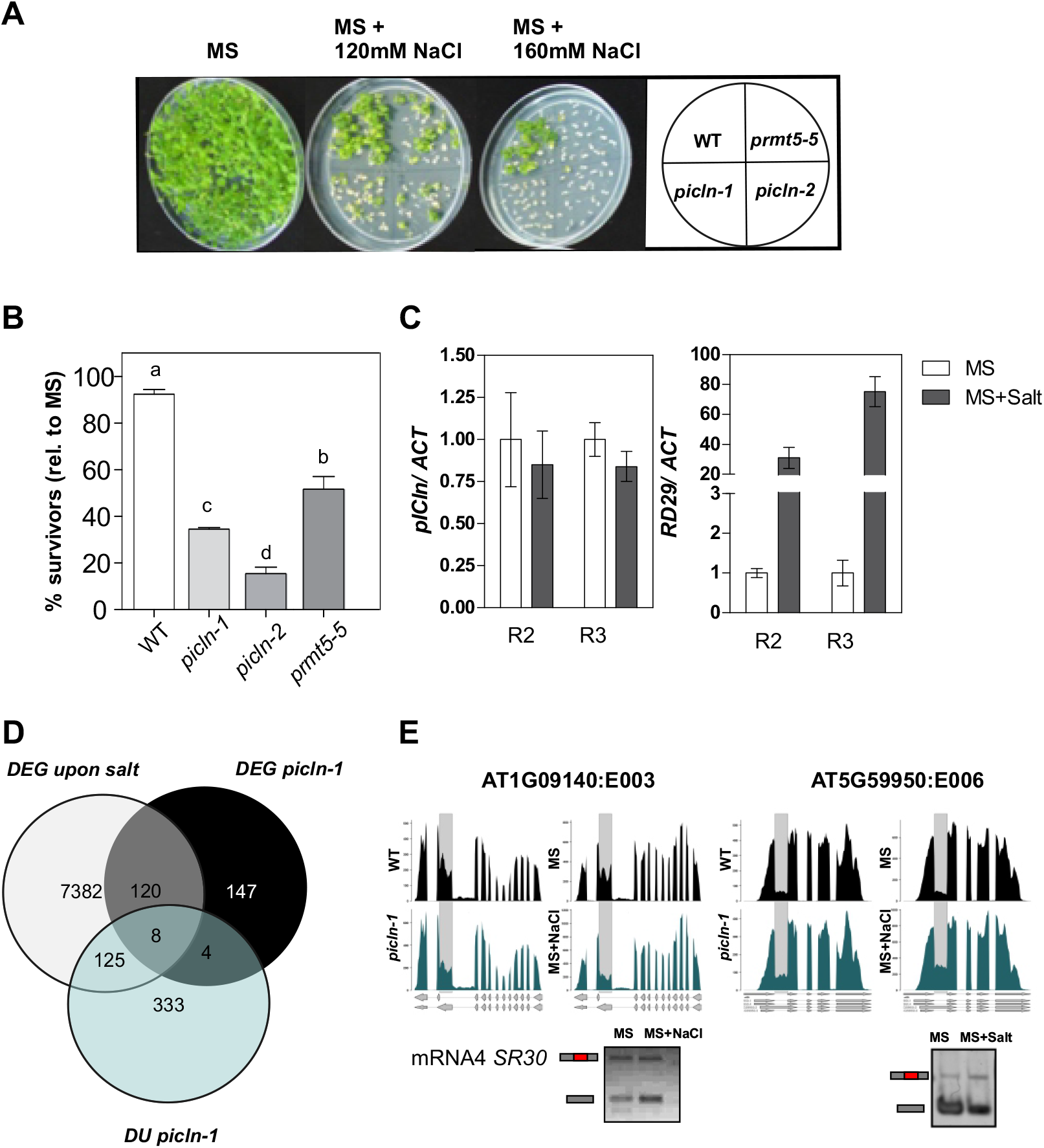
*pICln* salt sensitivity correlates with defects on alternative splicing of salt-responsive splicing factors. (A-B) *picln-1* mutant was hypersensitive to NaCl compared with wild-type plants in terms of survival. Survival rate on MS medium with 160 mM NaCl of wild-type plants and *picln*-1 mutants relative to controls without NaCl is shown. Survival was evaluated one month after seedlings were transferred to treatment medium. Letters indicate significant difference. (C) Root growth on MS medium with 100 mM NaCl. Primary root length of wild type and picln-1 mutants relative to controls grown without NaCl was determined 10 days after seedlings were transferred to treatment medium (n=12, *** =p.val <0.0001). (B) Expression level of pICln and RD29 (used as a salt responsive gene) in wild-type plants treated 7 h with 0 mM or 100 mM NaCl measured by qRT-PCR. Expression was normalized to ACTIN and shown relative to expression level in MS. Data are shown as mean ± SEM (n=3). Two biological replicates are shown (R2, R3). (C) Venn diagram showing the relationship among DEG upon salt treatment from Feng et. al 2015 and DEG and DU in picln from this study. Raw RNA-seq data from Feng et al. was downloaded and reanalyzed with the same pipeline used for picln datasets. (D) Browser with examples of two genes involved in splicing that respond to salt-treatment and are defective in picln.

## DISCUSSION

That Arabidopsis *picln* mutants, in contrast to corresponding mutants in many other eukaryotes, are viable has allowed for a detailed characterization of plant *PICLN* function during postembryonic development. *PICLN* has no paralogs in Arabidopsis, indicating that the viability of *picln* mutants is not due to genetic redundancy. Our characterization of *picln* mutants in Arabidopsis indicates that PICLN is a key modulator of plant growth and development in response to specific environmental perturbations rather than an essential gene.

Animal PICLN is a component of the methylosome complex, interacting with the SMN complex to modulate SnRNP assembly (Chari et al., 2008). The genome-wide analysis of splicing patterns conducted in *picln* mutants in this study establishes that most splicing events, particularly those constitutively processed in wild-type plants, do not display significant alterations in *picln* mutants. However, a subset of genes, most of which undergo AS in wild-type plants, showed alterations in the relative abundance of specific mRNA isoforms (Figure 3). Thus, PICLN is a modulator of AS, but its presence is not required for the completion of most splicing reactions.

PRMT5 and GEMIN2 are key components of the Methylosome and SMN complex, respectively, and are conserved from plants to humans. We found *picln* mutants to have many similarities with Arabidopsis *prmt5* and *gemin2* mutants. Indeed, mutations in the plant *PRMT5* or *GEMIN2* gene do not result in lethality, and their respective mutants are defective in specific subsets of AS events associated with adaptive responses to environmental conditions. These findings strongly suggest that PICLN, similarly to PRMT5 and GEMIN2, is important for modulating snRNP assembly and snRNP function in response to specific environmental perturbations, but less important for regular snRNP assembly under standard growth conditions. Consistent with this idea, *picln* mutants displayed very specific, rather than general, phenotypic alterations. These phenotypes included hypersensitivity to red, but not blue light; delayed flowering under intermediate, but not under long or short photoperiods; and growth impairment under cold conditions, but not under warmer temperatures as well as salt sensitivity (Figures 1 and 5-6).

PICLN physically interacts with PRMT5 in animals (Pu et al., 1999). Recently, Huang and colleagues (2016) found that Arabidopsis PRMT5 and PICLN interact *in vivo*, as determined by co-immunoprecipitation assays in *Nicotiana benthamiana* (Huang et al., 2016). Here, we have found an interaction between PICLN and PRMT5 by Y2H, strengthening the idea that these proteins form part of the same protein complex. Furthermore, we also observed interactions between PRMT5 and Sm proteins by Y2H, as well as between PICLN and Sm proteins (Figure 4), lending support to the idea that in plants PICLN also operates as a protein that assists in snRNPs assembly. Moreover, all Sm proteins that interact with PICLN were symmetrically dimethylated *in vivo* (Figure 4). Notably, unlike its human counterpart (Friesen et al., 2001a), PICLN does not appear to be required for PRMT5 methyl transferase activity in Arabidopsis (Figure 4). Thus, the molecular role that PICLN plays in association with the PRMT5 methylosome complex in plants remains to be determined.

In addition to the physical interaction detected between PICLN and PRMT5, the expression patterns of *PICLN* and *PRMT5* genes were strongly correlated, further suggesting that the proteins they encode act in part together to control AS. While we obtained some support for this hypothesis, as indeed the *picln* and *prmt5* mutants share many molecular and physiological phenotypes, they also displayed several distinct physiological and molecular alterations, indicating that PICLN and PRMT5 have both overlapping and distinct molecular functions and regulatory roles. For example, although the *PICLN* gene is clock-regulated at the mRNA level, *picln* mutants do not affect circadian rhythmicity, in contrast to *prmt5* mutants (Sanchez et al., 2010). This observation suggests that circadian regulation of *PICLN* expression contributes to the control of clock output pathways but not to the control of the core oscillator itself. Thus, in relation to the circadian gene network, the role of PICLN is more similar to that we recently reported for the Arabidopsis *Survival Motor Neuron-like* gene *SPLICING FACTOR 30* (*SPF30*), whose expression is clock-regulated and its function modulates circadian oscillations for a subset of AS events but does not affect circadian rhythms in general (Romanowski et al., 2020).

The changes in AS patterns in *picln* mutants resemble those observed in *gemin2* mutants, and less so those seen in *prmt5* mutants (Figure 3, Supplemental Figure S6). The functional isoform of U1-70K was the predominant isoform in *picln* and *gemin2* mutants, while the version retaining an intron was more abundant in wild type and *prmt5* mutants. Also, retention of intron 2 of *SPF30* was more pronounced in the wild type than in *picln* or *gemin2* mutants (Supplemental Figure 7C), reinforcing the parallels between these genotypes. Similarities are not only associated with the molecular phenotypes of these mutants, but also with the regulation of their expression in response to environmental signals. Indeed, as previously observed for *GEMIN2, PICLN* expression was also induced in response to low temperatures (Figure 5, Figure S12), a condition in which PICLN as well as GEMIN2 proteins played a key role in modulating AS. Thus, regulation of *PICLN* expression is coordinated with PICLN function to ensure the optimal operation, i.e attenuating the deleterious effects of abiotic stress, particularly cold conditions, on AS, and, as a consequence, on plant growth and proper development. Likewise, GEMIN2 is also required for proper growth and survival under cold conditions. Indeed, *gemin2* mutants do not survive after two weeks at 4°C (Schlaen et al., 2015).

We propose that PICLN, similarly to PRMT5 and GEMIN2, is a key modulator of AS in response to specific environmental perturbations or endogenous signals, and the expression of its encoding gene is modulated by these same environmental or endogenous signals to ensure optimal control of splicing reactions throughout the day and year. Indeed, we also observed that *SmB, SmD2, SmD3, SmE1*/*PCP* and *SmG* are upregulated upon cold treatments (Supplemental Table S1), likely to compensate for the negative effect of temperature on snRNP function and thus maintain proper splicing.

PICLN is an unusual protein in that it exerts multiple functions in different complexes and, therefore, influences multiple biological processes. This mode of action could be the result of its possible operation as a chaperone or adaptor protein that may modulate the assembly of different macromolecular complexes. In the early phase of snRNP assembly, PICLN mimics the Sm fold, and enables Sm proteins to interact with each other even in the absence of the snRNA, thus allowing complex formation. This ability relies in part on the Pleckstrin homology (PH) domain at the C-terminal region of the protein, which is mostly an unstructured domain that becomes ordered upon formation of the snRNP complex (Grimm et al., 2013).

Why PICLN is not essential in plants and yeast but is required for viability in animals is not clear (Pu et al., 2000; Barbarossa et al., 2014). It is unlikely to be essential for spliceosome assembly in plants but not animal cells, considering the similarities between spliceosomal components. Functional redundancy with other genes is excluded. In addition, the rescue of the Arabidopsis *picln* mutant by the human PICLN ortholog gene strongly suggests that the molecular functions of animal and plant proteins are conserved. Perhaps PICLN is not essential for viability of all animal cells, but for a subset or subtype, with particular cellular environments in which subtle alterations in splicing due to delay or suboptimal snRNP assembly leads to cell death, and eventually to organismal malfunction. The role of PICLN in different individual animal cells needs to be addressed to evaluate this idea in detail.

Splicing is a complex process involving multiple RNA and protein components. Spliceosome assembly and splicing reactions involve many interactions and rearrangements that require coordination and assistance. Although the assembly can take place autocatalytically, without requiring specific assembly factors, stress conditions challenge this situation; it is under these challenging conditions that PICLN, PRMT5 and GEMIN2 become critical players to ensure homeostatic operation of splicing and maximize survival, growth and development. As for many other splicing-related proteins, it remains to be determined why PICLN plays a key role for the proper splicing of only a subset of genes.

## Methods

### Plant material and growth conditions

All Arabidopsis lines used in this study were in the Columbia-0 (Col-0) accession. Plants were grown on soil under controlled conditions at 22°C under different light regimes depending on the experiment. The *picln-1* (GABI_675D10) and *picln-2* (SALK_050231) mutants were obtained from the ABRC and verified by PCR. For genetic analysis, *prmt5-5* (Sanchez et al., 2010), *gemin2-1* (SALK_142993) (Schlaen et al., 2015), *smd1a-2* (GABI_380A07), *sme1-1* (GABI_207E07), *smd3b* (SALK_006410C) (Swaraz et al., 2011) were crossed to *picln-1*. In the F_2_ and F_3_ generations, plants were genotyped with the primers listed in Supplemental Table S2.

For RNA-seq experiments, *picln-1, prmt5-5* and wild-type seedlings were grown under 12-h light/12-h dark conditions. Four biological replicates were collected. Seeds were sown onto Murashige and Skoog (MS) medium containing 0.8% (w/v) agar and were stratified for 4 d in the dark at 4°C, before being released at 22°C in a 12-h light/12-h dark cycle. Whole seedlings were harvested after 9 d and total RNA was extracted with RNeasy Plant Mini Kit (QIAGEN) following the manufacturer’s protocols. To estimate concentration and quality of all samples, NanoDrop 2000c (Thermo Scientific) and the Agilent 2100 Bioanalyzer (Agilent Technologies) with the Agilent RNA 6000 NanoKit were used, respectively. Libraries were prepared following the TruSeq RNA Sample Preparation Guide (Illumina). Samples were pooled to create 12 multiplexed libraries, which were sequenced as single-end reads with an Illumina Genome Analyzer II Kit on the Illumina GAIIx platform, providing 100-bp reads.

For RNA-seq of the cold treatment experiment, seeds from three biological replicates were sown onto MS medium containing 0.8% (w/v) agarose, stratified for 3 d in the dark at 4°C and then grown at 22°C for 9 d under continuous light. On day 10, plates were transferred to 10°C or kept at 22°C for 24 h before sample collection. Total RNA was extracted with TRIZOL or Spectrum RNA extraction Kit (Sigma). Illumina libraries were prepared at Macrogen Korea and 150-bp reads (pair-end) were obtained after high-throughput sequencing.

### Phenotypic analysis

For determination of flowering phenotypes, the number of rosette leaves were counted at flowering (when bolting) at 22°C for at least 12 individual plants under three photoperiods (long-day:16-h light/8-h, intermediate:12h light/12-h, or short-day: 8-h light/16-h). For leaf movement assays, seedlings were grown at 22°C under 16-h light/8-h dark cycles for 10 d, transferred to continuous white fluorescent light (at 22°C or 17°C) and the position of the first pair of true leaves was recorded every hour for 6 d using digital cameras. Leaf angle was determined using ImageJ software. Period estimates were calculated with Brass 3.0 software (Biological Rhythms Analysis Software System, available from http://www.amillar.org) and analyzed with the FFT-NLLS suite of programs. For hypocotyl length measurements, seeds were sown in 0.8% (w/v) water agar and stratified for 4 d at 4°C in the dark. Germination was induced by a 2-h white light pulse, followed by 22 h of darkness. Seedlings were then grown in darkness, continuous red or continuous blue light and hypocotyl length was measured after 3 d (six replicates of 10 seedlings each). For salt stress assays, seeds were germinated on MS agar medium and 5-d-old seedlings grown under 16-h light/8-h dark cycles were transferred to MS agar medium containing different concentrations of NaCl. For primary root length experiments, root length was measured 10 d after transferring seedlings to 0 or 100 mM NaCl. To score the embryo-lethality phenotype of double mutants, selfed siliques from segregating F_3_ plants (with a single wild-type copy of either gene) as well as those of parental single mutants were dissected and photographed with a Leica DMS1000 macroscope. Both phenotypically wild-type and abortive seeds were classified and cleared in a mix of chloral hydrate:glycerol:water (8:1:3, v/v/v) for 2 to 6 weeks and then observed under differential contrast microscope (DIC) optics in a Nikon Eclipse E600 microscope.

### Plasmid construction

To generate heterologous expression constructs, the coding sequences of *PICLN* obtained from human or Arabidopsis were cloned into the pEarleyGate 100 and pB7WYF2 vectors, respectively, and introduced into Agrobacterium (*Agrobacterium tumefaciens GV3101*). *picln-1* mutant plants were transformed by the floral dip method.

The coding sequences of the *SmB* (At4g20440), *SmD1A* (At3g07590) and *SmD3* (At1g76300) genes were obtained from the Arabidopsis Information Resource (TAIR10), synthesized de novo by GenScript Corporation, and cloned into the pB7WYF2, pB7WCF2 and pB7WGF2 destination vectors, respectively, using Gateway technology (Invitrogen). These constructs were used for immunoprecipitation experiments.

For yeast two-hybrid constructs involving interactions between PICLN and Sm proteins, pENTR223 clones containing the *PICLN, SmBb, SmD1A, SmD2A, SmD3B, SmE1/PCP* and *SmF* coding sequences were obtained from the Arabidopsis Biological Resource Center. The *SmGb* coding sequence was amplified from a pool of Arabidopsis cDNA and cloned into the pDONR221 via BP reaction (Invitrogen). For yeast two-hybrid constructs involving interactions with PRMT5 coding sequences of the genes of interest were cloned into pGADT7 or pGBKT7 vectors.

### RNA extraction and quantitative real-time PCR

Total RNA was isolated from seedlings using TRIZOL (Ambion) and treated with RNAse-free DNase I (Promega) to remove residual genomic DNA. One microgram of total RNA was used for reverse transcription (Superscript II, Invitrogen). Transcript levels were quantified by quantitative PCR in a Stratagene MX3005P instrument (Agilent Technologies) using *PP2A* (At1g13320) or *ACTIN* (At5g09810) as housekeeping gene. The sequences of the primers used to quantify the expression are listed in Supplemental Methods.

### Validation of splicing events

cDNAs was synthesized as above but with SuperScriptIII (Thermofischer). PCR amplification was performed using 1.5 U of Taq polymerase (Invitrogen). Primers used for amplification are detailed in Supplemental Table S2. RT-PCR products were electrophoresed on 2% (w/v) agarose gels and stained with SYBR Green or ethidium bromide. The band corresponding to different isoforms were quantified by image analysis and ratios were reported. We selected the events to validate according to the following parameters: the gene should have at least 50 reads to ensure good expression, the difference between the isoforms should not be greater than 10% of the size and when possible, the neighbor intron without defect should be included in the PCR product to show that the event we measure is specific.

### Yeast two-hybrid assays

For Yeast two-hybrid involving interactions with PICLN, assays were conducted after mating Y187 and Y2H Gold haploid strains transformed with various GAL4-activation domain and GAL4-DNA binding domain fusions, respectively. Interactions were assayed on synthetic define (SD) medium lacking histidine. To raise sensitivity, 5 mM 3-amino-1,2,4-triazole (3-AT) was added. For Y2H assays involving PRMT5 pairs of vectors were used to co-transform AH109 yeast strain and then positive colonies carrying both vectors were selected on SD-WL at 28°C. After 6 days, protein-protein interactions were tested by growing serial dilutions also on SD-WLH media.

### Protein analysis

Total protein extraction and *in vivo* immunoprecipitation (IP) were performed as described previously with minor modifications (Esteve-Bruna et al., 2020). IP was performed using anti-GFP beads (Chromotek-gta-20) or anti-GFP-coated paramagnetic beads (Miltenyi) following the manufacturer’s instructions. For symmetric dimethylation detection, immunoblot analysis was performed with the anti-dimethyl-arginine antibody, symmetric (SYM10). For GFP detection, anti-GFP from Abcam (ab290) or Clontech (JL8) were used. Roughly 50 15-day-old LD-grown seedlings were harvested.

Protein concentration was determined with a Bradford assay and a total of 50 μg for crude extracts were separated for immunoblots or 1 mg was used for immunoprecipitations.

### Alternative splicing analysis

Raw reads generated from this study were mapped to the Arabidopsis genome (TAIR10) using the TopHat-Bowtie pipeline (Trapnell et al., 2009) with default parameters except for maximum intron length set to 5,000 nucleotides. Differential gene expression analysis and differential splicing were analyzed using R with the ASpli package (Mancini et al., 2021). For expression analysis, genes with a false-discovery rate (FDR)<0.05 were considered as differentially expressed between genotypes. For splicing analysis, PSI (percent of inclusion) and PIR (percent of IR) were calculated and differential splicing was considered for bins with FDR <0.15 and ΔPSI/PIR >0.05.

For the analyses of RNA-seq data from salt treatment, raw reads from GSE55632 (Feng et al., 2015) were downloaded from NCBI and treated as described above.

## Data availability

The RNA-seq data generated in this study have been deposited at the GEO under the accession number GSE130463 (for 12-h light/12-h dark treatment) and in SRA for cold treatments experiments under the number PRJNA559541.

## Acknowledgments

We thank Prof. Dorothee Staiger and Dr. Marlene Reichel for critical reading of the manuscript.

## Author Contributions

SES, JLM and MJY designed the research, SS, ML, DE, DG, DKS, NB-T, JT, EP, JLM performed the experiments, JLM and MJY wrote the paper with contributions from all the authors.

## SUPPLEMENTS

**Supplemental Figure S1. *PICLN* expression is regulated by the circadian clock but it does not have a role in circadian leaf movements**.

(A) Leaf number at flowering time estimated under long-day (16-h light/8-h dark, left) or short-day (8-h light/16-h dark, right) conditions. Significant differences were determined by Tukey’s test for multiple comparisons (n>18 plants, p<0.0001). (B) (Top) *PICLN* transcript levels show a circadian rhythm (At5g62290). Seedlings were entrained under a long-day photoperiod (16-h light/8-h dark) before being transferred to constant light (LL). Expression is shown for d 2 and 3 in LL after light-dark entrainment. Data from http://diurnal.mocklerlab.org/. (Bottom) Leaf movement traces for the indicated genotypes. Leaf angles were measured for the first pair of leaves in seedlings first entrained under long-day conditions (16-h light/8-h darkness) before being transferred to constant light (LL). Wild-type (Col-0) (open boxes; n=3), *picln-1* (light-gray triangles; n=6), *picln-2* (dark-gray triangles; n=6). (C) Pictures of *picln-1* and *picln-2* compared to wild-type showing reduced growth. Plants were grown in long-day photoperiod. (D) Leaves from wild-type and *picln-1*. All rosette leaves analyzed until flowering in long-day photoperiod.

**Supplemental Figure S2. Expression levels of components of the splicing machinery in *picln* mutants**

(A) Expression of *SMs, GEMIN2* and *PRMT5* transcripts in 12-d-old seedlings from wild-type or *picln-1* grown under 12L:12D photoperiod at 22°C. (B) Expression levels of *PICLN* in *gemin2-1* and *prmt5-5* mutants. Data was obtained from RNA-seq experiments. (C) Selection of Gene Ontology-terms enriched among differentially expressed genes in *picln-1* in comparison to wild-type plants.

**Supplemental Figure S3. Alleles used in this study**

(A) Gene model of *SMD1A* (At3g07590) and *SMEB* (At2g18740). Black boxes, exons; white boxes, UTRs; black lines, introns. T-DNA insertion sites in *smd1a-1* and *sme1-1* mutants are indicated with white triangles. Primers to measure transcript levels are shown below the gene model as black arrows. (B) *SMD1A* expression in wild-type and *smd1a-1* or *SME1B* expression in wild-type and *smeb-1* seedlings, as measured by end-point RT-PCR. *UBQ10* was used as a control.

**Supplemental Figure S4. Genetic interaction between *PICLN* and several *Sm* genes**

(A-E) Phenotypes of *picln-1* (A), *smd1a-1* (B) and *picln-1 smd1a-1* (C) 14-day-old mutant plants. (F-G) Developing seeds from siliques of selfed *picln-1* (F) or *picln-1/picln-1 SMD3B/smd3b* (G) plants. Arrows indicate aborted double mutant seeds. (H-J) DIC images of seeds from homozygous or segregating genotypes, the *picln-1* single mutant (H) and the *picln-1 smd3b* double mutant. Note that embryo development in arrested at the torpedo stage in the double mutant (I-J).

**Supplemental Figure S5. Validation of splicing events affected in *picln-1***.

(Top) Two examples of splicing events affected in *picln-1*, visualized as coverage plots from RNA-seq experiments. “I006” and “E008” defines the sixth intronic bin or eight exonic bin, respectively. (Bottom) Confirmation by RT-PCR of the defects shown at the top. Alternative regions are highlighted in red in the diagrams next to the gels.

**Supplemental Figure S6. Transcriptome of *picln-1* compared to *prmt5-5, sme1/pcp-1 and gemin2-1***.

(A-C) Left: Venn diagram showing the extent of overlap for pre-mRNA splicing events affected in the *picIn-1* mutant and those altered in *prmt5-5* (A), *sme1-1/pcp-1* (B), and *gemin2-1* (C). Middle: Venn diagram showing the extent of overlap between differentially expressed genes in *picln-1* and those in *prmt5-5* (A), *sme1-1/pcp-1* (B) and *gemin2-1* (C). Right: Hierarchical clustering of common differentially expressed genes between *picln-1* and either *prmt5-5, pcp-1* or *gemin2-1*. Clustering was done with the complete linkage method with Clustal 3.0 Software. Values represent Log2FC among genotypes. Red arrow: upregulated genes; green arrow: downregulated genes

**Supplemental Figure S6. Examples of selected splicing defects in *picln-1* compared to *prmt5-5, sme1 and gemin2***.

(A-C) Integrated genome browser view showing coverage plots from nine selected genes with splicing defects in *picln-1* and/or *prmt5-5* (A), *sme1-1/pcp-1* (B) or *gemin2-1* (C). Data for *picln-1* and *prmt5-5* were obtained in this study, from seedlings grown for 12 d in 12-h light/12-h dark conditions. Data for *gemin2-1* were obtained from (Schlaen et al., 2015); data for *pcp-1* from (Capovilla et al., 2018). (D) Read density map of U1-70K in each mutant, as visualized in IGV. The alternatively spliced intron is highlighted with a box.

**Supplemental Figure S8. Confirmation of splicing defects in *picln-1* compared to *prmt5-5***

(Left) Integrated genome browser view showing coverage plots from three genes with splicing defects in *picln-1* compared to wild type and *prmt5-5*. (Right) Confirmation by RT-PCR of the defects shown in the left. Alternative regions are highlighted in red in the diagrams next to the gels.

**Supplemental Figure S9. Sequences of Arabidopsis Sm proteins**

Protein sequences of Arabidopsis Sm proteins. RG motifs are highlighted in red.

**Supplemental Figure S10. PRMT5 can interact with Sm proteins and PICLN in yeast**.

(A) Interactions between Sm proteins and PRMT5 or PRMT5 and PICLN are shown. AD and BD domains are carried by pGADT7 and pGBKT7 vector, respectively. After transformation, yeast was grown on -WL selective media for 6 days at 28°C, later serial dilutions of the resulting colonies were grown on -WL and -WLH selective media for additional 4 days before taking pictures. This panel represents the full content of Figure 4B. (B) negative controls of the experiment in A are shown.

**Supplemental Figure S11. Subcellular localization of SmB, SmD1, SmD3 and PICLN**.

(A) Subcellular localization of SmB:YFP, SmD1:CFP and SmD3:GFP under the control of the 35S CaMV promotor in wild-type or *picln-1* background. (B) Subcellular localization of PICLN::YFP. Pictures we taken from hypocotyl cells (a) or root cells (b). Arrows indicate the cytoplasm while arrow heads indicate nuclei. Middle panel represents DAPI staining and right panel Merge between YFP channel and DAPI.

**Supplemental Figure S12. Expression levels of components of the splicing machinery under cold**.

(A) Cold sensitivity of wild type and *picln-1* at 10°C. Four-day-old seedlings grown at 22°C in a 16-h light/8-h dark photoperiod were kept at 22°C as control or transferred to continuous light at 10°C. Pictures were taken 3 weeks or 8 weeks later, respectively. (B) Expression of *SMs, GEMIN2* and *PRMT5* transcripts in 10-days-old seedlings grown at 22°C or grown at 22°C and exposed to 10°C for 24 hours.

### Supplemental Tables

**Supplemental Table S1: RNA-seq from *picln*-1 12-h light/12-h dark** (showing DEG and DU), and **RNA-seq from *picln*-1 in cold**: (showing DEG and DU). Several comparisons are described.

**Supplemental Table S2: Primers**

