## Supplemental Figures for "PICLN modulates alternative splicing and ensures adaptation to light and temperature changes in plants"

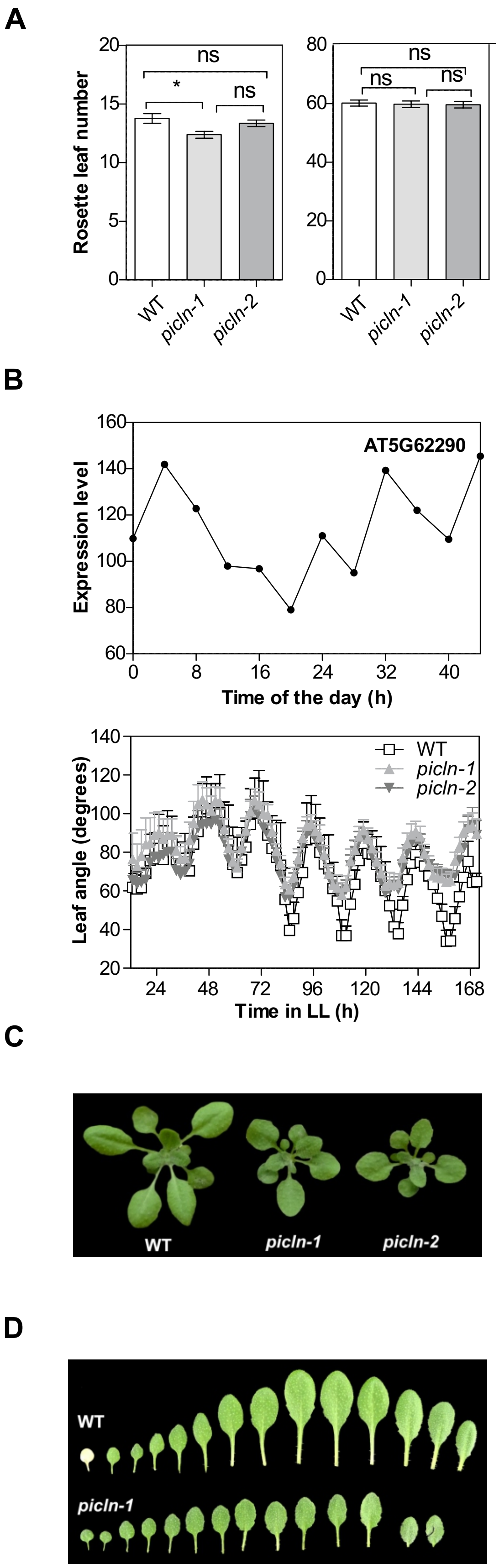
A
Rosette leaf number
ns
*
ns
ns
ns
ns
WT
picln-1
picln-2
B
AT5G62290
Expression level
Time of the day (h)
Leaf angle (degrees)
WT
picln-1
picln-2
Time in LL (h)
C
WT
picln-1
picln-2
D
WT
picln-1

**Figure S1**

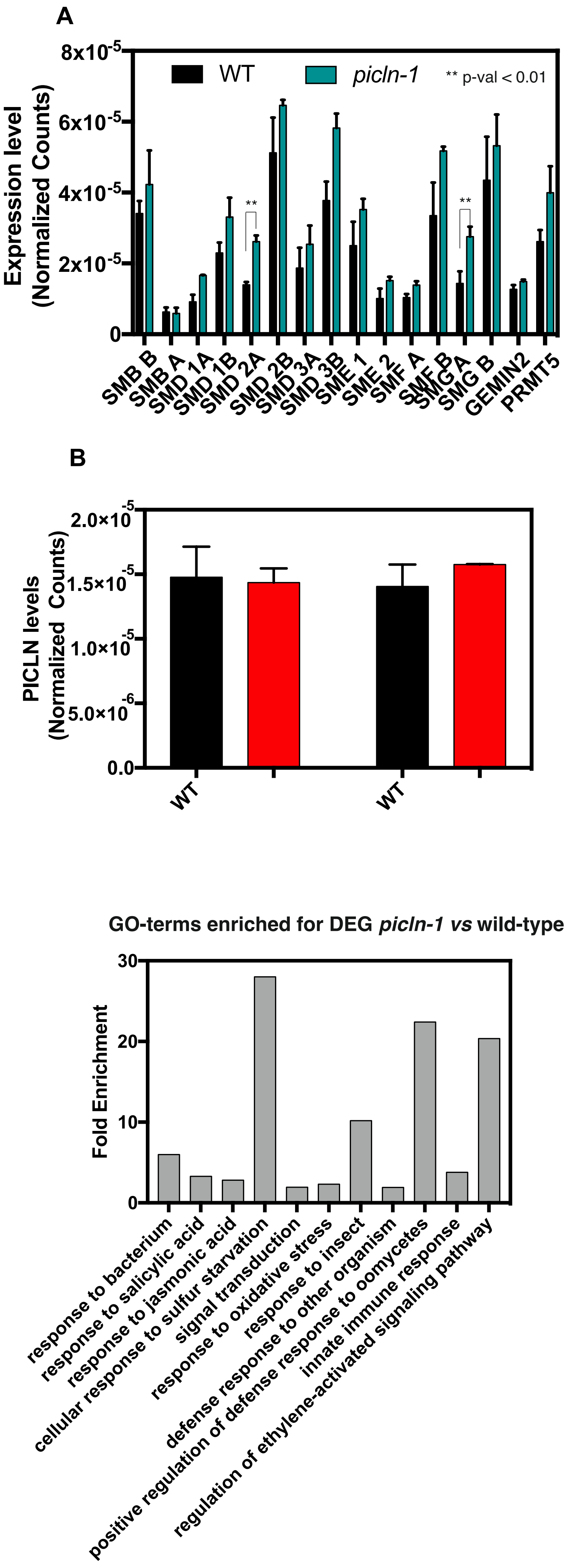
A
8x10-5
6x10-5
4x10-5
2x10-5
0
Expression level
(Normalized Counts)
WT
picln-1
** p-val < 0.01
**
**
SMB B
SMB A
SMD 1A
SMD 1B
SMD 2A
SMD 2B
SMD 3A
SMD 3B
SME 1
SME 2
SMF A
SMF B
SMG A
SMG B
GEMIN2
PRMT5
B
2.0×10-5
1.5×10-5
1.0×10-5
5.0×10-6
0.0
PICLN levels
(Normalized Counts)
WT
WT
GO-terms enriched for DEG picln-1 vs wild-type
30
20
10
0
Fold Enrichment
response to bacterium
response to salicylic acid
response to jasmonic acid
cellular response to sulfur starvation
signal transduction
response to oxidative stress
response to insect
defense response to other organism
positive regulation of defense response to oomycetes
innate immune response
regulation of ethylene-activated signaling pathway

**Figure S2**

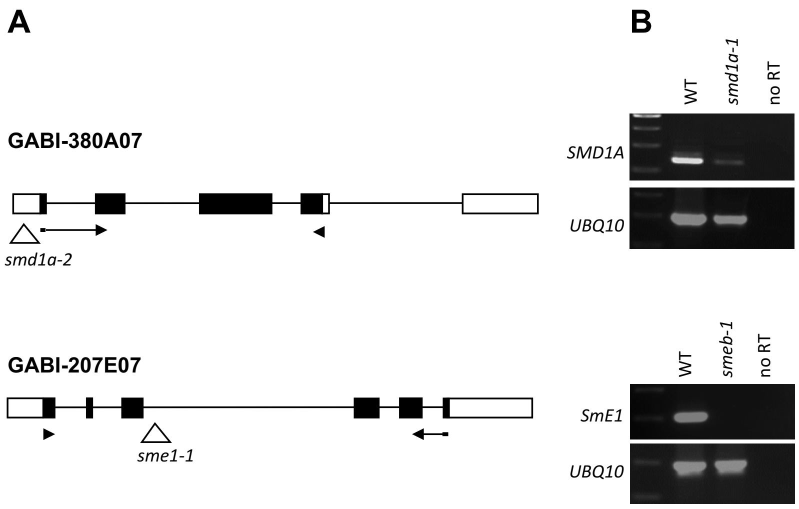
A
GABI-380A07
smd1a-2
GABI-207E07
sme1-1
B
WT
smd1a-1
no RT
SMD1A
UBQ10
WT
smeb-1
no RT
SmE1
UBQ10

**Figure S3**

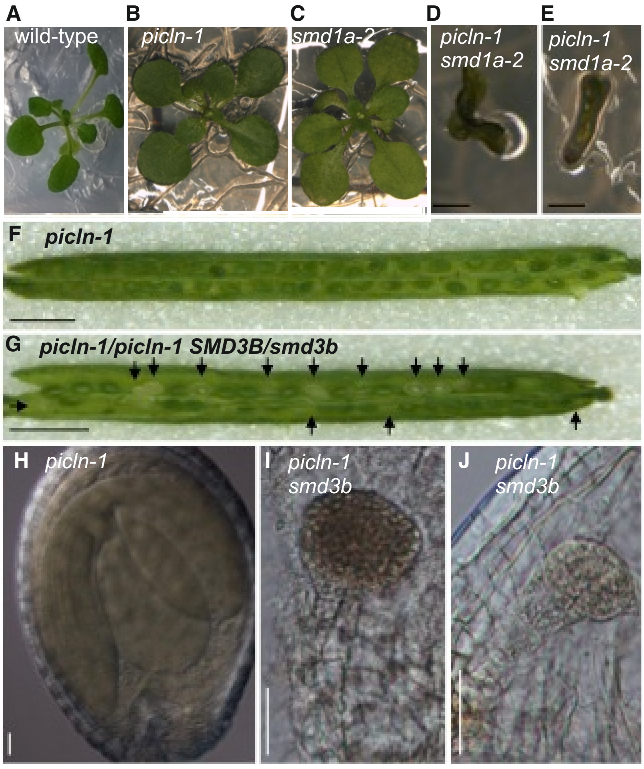
A
wild-type
B
picln-1
C
smd1a-2
D
picln-1
smd1a-2
E
picln-1
smd1a-2
F
picln-1
G
picln-1/picln-1 SMD3B/smd3b
H
picln-1
I
picln-1
smd3b
J
picln-1
smd3b

**Figure S4**

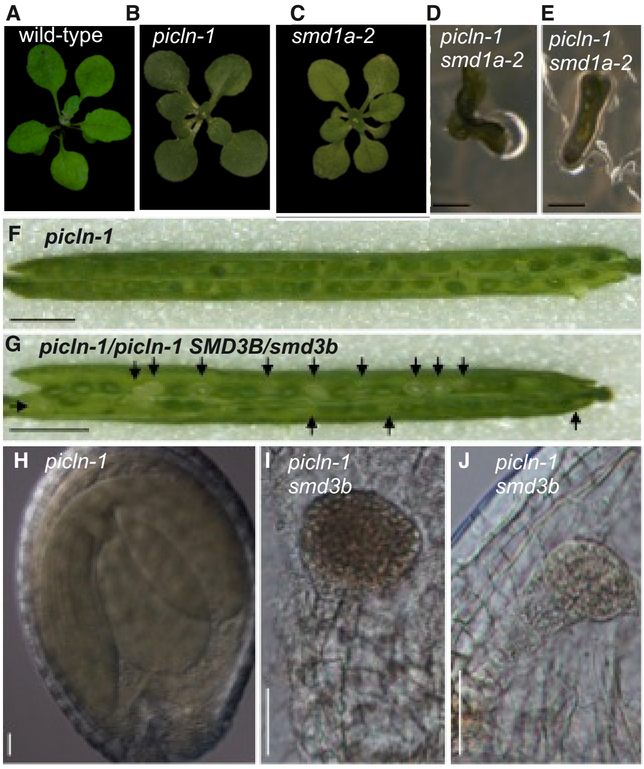
A
wild-type
B
picln-1
C
smd1a-2
D
picln-1
smd1a-2
E
picln-1
smd1a-2
F
picln-1
G
picln-1/picln-1 SMD3B/smd3b
H
picln-1
I
picln-1
smd3b
J
picln-1
smd3b

**Figure S4**

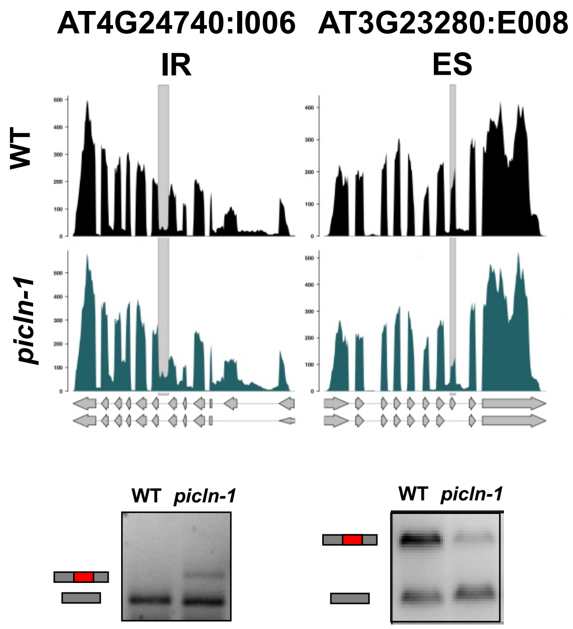

AT4G24740:I006
AT3G23280:E008
IR
ES
WT
picln-1
WT
picln-1
WT
picln-1


**Figure S5**

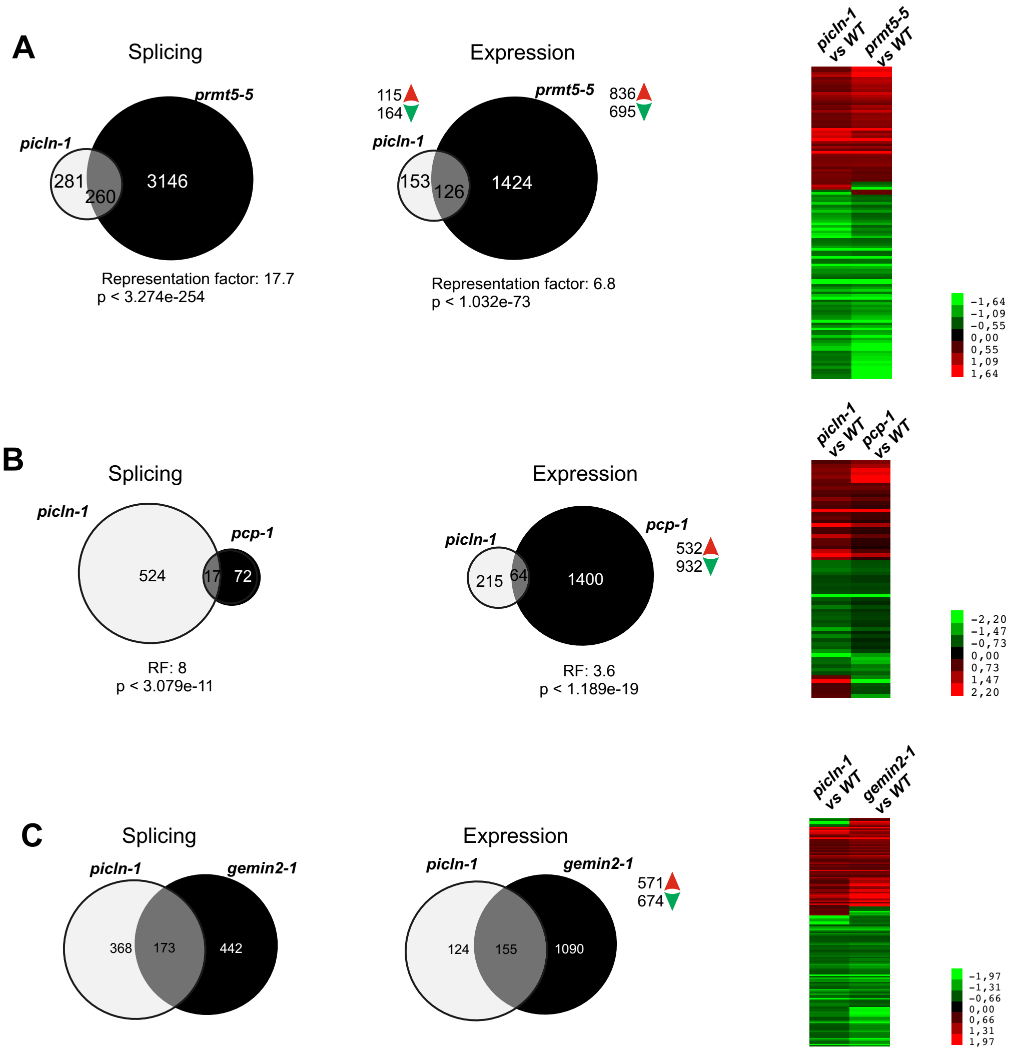
A
Splicing
picln-1
prmt5-5
281
260
3146
Representation factor: 17.7
p < 3.274e-254
Expression
115
164
836
695
153
126
1424
Representation factor: 6.8
p < 1.032e-73
picln-1 vs WT
prmt5-5 vs WT
-1,64
-1,09
-0,55
0,00
0,55
1,09
1,64
B
Splicing
pcp-1
524
17
72
RF: 8
p < 3.079e-11
Expression
215
64
1400
532
932
RF: 3.6
p < 1.189e-19
pcp-1 vs WT
-2,20
-1,47
-0,73
0,00
0,73
1,47
2,20
C
Splicing
gemin2-1
368
173
442
Expression
124
155
1090
571
674
gemin2-1 vs WT
-1,97
-1,31
-0,66
0,00
0,66
1,31
1,97

**Figure S6**

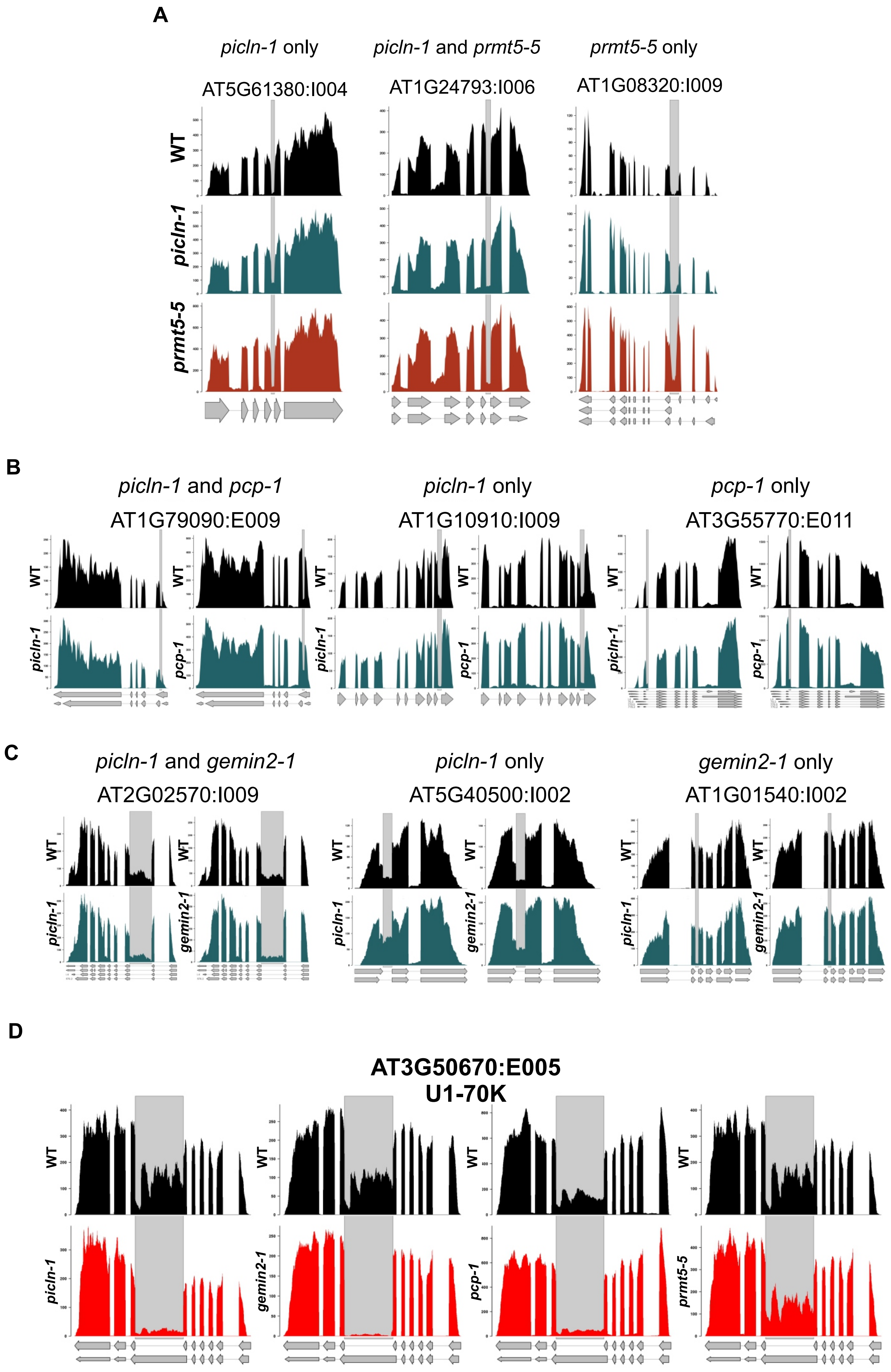
A
picln-1 only
picln-1 and prmt5-5
prmt5-5 only
AT5G61380:I004
AT1G24793:I006
AT1G08320:I009
WT
picln-1
prmt5-5
B
picln-1 and pcp-1
picln-1 only
pcp-1 only
AT1G79090:E009
AT1G10910:I009
AT3G55770:E011
WT
picln-1
pcp-1
C
picln-1 and gemin2-1
picln-1 only
gemin2-1 only
AT2G02570:I009
AT5G40500:I002
AT1G01540:I002
WT
picln-1
gemin2-1
D
AT3G50670:E005
U1-70K
WT
picln-1
gemin2-1
pcp-1
prmt5-5

**Figure S7**

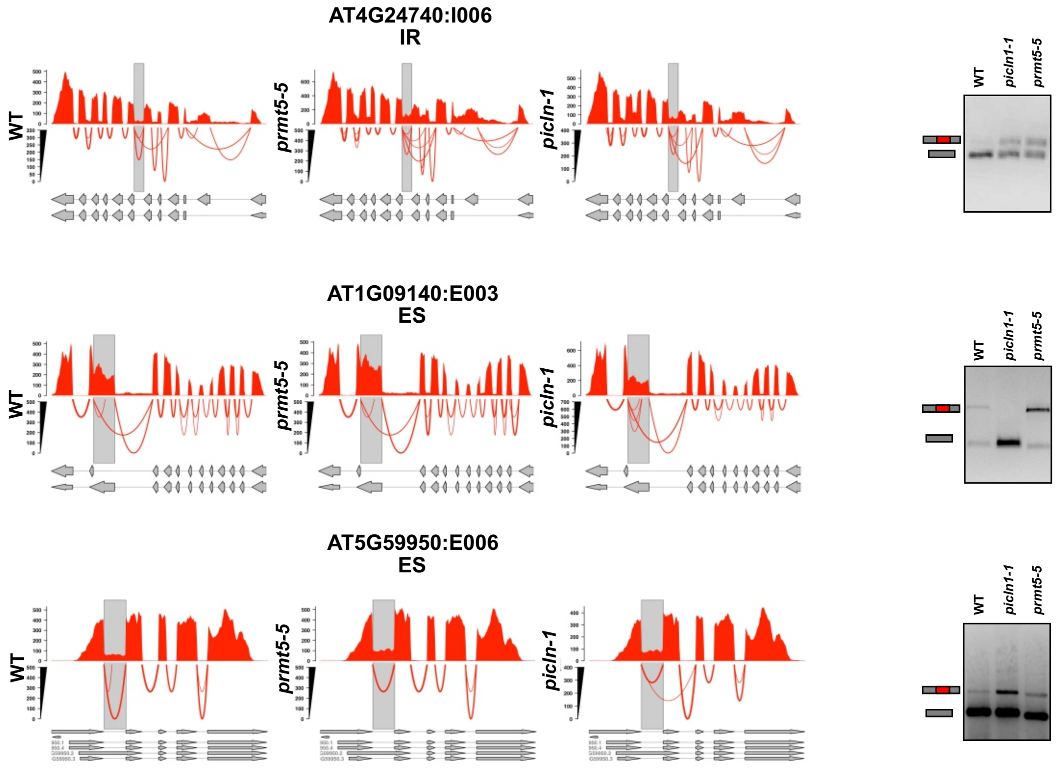
AT4G24740:I006
IR
WT
prmt5-5
picln-1
picln1-1
AT1G09140:E003
ES
AT5G59950:E006
ES

**Figure S8**

```
>SMBA
  1 MSMSKSSKML QFINYRMRVT IQDGRQLIGK FMAFDRHMNL VLGDCEEFRK
 51 LPPAKGNKKT NEEREERRTL GLVLLRGEEV ISMTVEGPPP PEESRAKSGS
101 VTAVAGPGIG RAAGRGVPTG PLVQAQPGLS GPVRGIGGPA PGMMQPQISR
151 PPQIIRPPGQ MPPQPPFAGQ GGPPPPYGMR PPYPGPPPPQ YGGQQRPMMI
201 PPPGGMMRGP PPPHGMQGPP PSRPGMPPPG GAPMFAPPHP GMPPAPPNHH
251 NQQH

>SMBB
  1 MSMSKSSKML QFINYRMRVT IQDGRQLVGK FMAFDRHMNL VLGDCEEFRK
 51 LPPAKGKKIN EEREDRRTLG LVLLRGEEVI SMTVEGPPPP EESRAKAGSA
101 AAVAGPGIGR AAGRGVPTGP LVQAQPGLSG PVRGVGGPAP GMMQPQISRP
151 PQLSAPPIIR PPGQMLPPPP FGGQGPPMGR GPPPPYGMRP PPQQFSGPPP
201 PQYGQRPMIP PPGGMMRGPP PPPHGMQGPP PPRPGMPPAP GGFAPPRPGM
251 PPHNQQQ

>SMD1A
  1 MKLVRFLMKL NNETVSIELK NGTVVHGTIT GVDVSMNTHL KTVKMSLKGK
 51 NPVTLDHLSL RGNNIRYYIL PDSLNLETLL VEDTPRVKPK KPVAGKAVGR
101 GRGRGRGRGR GRGR

>SMD1B
  1 MKLVRFLMKL NNETVSIELK NGTIVHGTIT DEYTRFYLNR GVDVSMNTHL
 51 KAVKLTLKGK NPVTLDHLSV RGNNIRYYIL PDSLNLETLL VEDTPRIKPK
101 KPTAGKIPAG RGRGRGRGRG RGRGGR

>SMD2A
  1 MSKPMEEDTN QGKTEEEEFN TGPLSVLMMS VKNNTQVLIN CRNNRKLLGR
 51 VRAFDRHCNM VLENVREMWT EVPKTGKGKK KALPVNRDRF ISKMFLRGDS
101 VIIVLRNPK

>SMD2B
  1 MSKPMEEDTN QGKTEEEEFN TGPLSVLMMS VKNNTQVLIN CRNNRKLLGR
 51 VRAFDRHCNM VLENVREMWT EVPKTGKGKK KALPVNRDRF ISKMFLRGDS
101 VIIVLRNPK

>SMD3A
  1 MSRSLGIPVK LLHESSGHIV SVEMKSGELY RGSMIECEDN WNCQLENITY
 51 TAKDGKVSQL EHVFIRGSLV RFLVIPDMLK NAPMFKDVRG KGKSASLGVG
101 RGRGAAMRAK GTGRGTGGGR GAVPPVRR

>SMD3B
  1 MSRSLGIPVK LLHEASGHIV TVELKSGELY RGSMIECEDN WNCQLEDITY
 51 TAKDGKVSQL EHVFIRGSKV RFMVIPDILK HAPMFKRLDA RIKGKSSSLG
101 VGRGRGAMRG KPAAGPGRGT GGRGAVPPVR R

>SME1
 1 MASTKVQRIM TQPINLIFRF LQSKARIQIW LFEQKDLRIE GRITGFDEYM
51 NLVLDEAEEV SIKKNTRKPL GRILLKGDNI TLMMNTGK

>SME2
 1 MASTKVQRIM TQPINLIFRF LQSKARIQIW LFEQKDLRIE GRITGFDEYM
51 NLVLDEAEEV SIKKKTRKPL GRILLKGDNI TLMMNAGK

>SMF
 1 MAFVCMCVLQ TIPVNPKPFL NNLTGKTVIV KLKWGMEYKG FLASVDSYMN
51 LQLGNTEEYI DGQLTGNLGE ILIRCNNVLY VRGVPEDEEL EDADQD

>SMGA
 1 MSRSGQPPDL KKYMDKKLQI KLNANRMVTG TLRGFDQFMN LVVDNTVEVN
51 GNDKTDIGMV VIRGNSIVTV EALEPVGRSS

>SMGB
1 MSRSGQPPDL KKYMDKKLQI KLNANRMVVG TLRGFDQFMN LVVDNTVEVN
51 GDDKTDIGMV VIRGNSIVTV EALEPVGRS
```

**Figure S9**

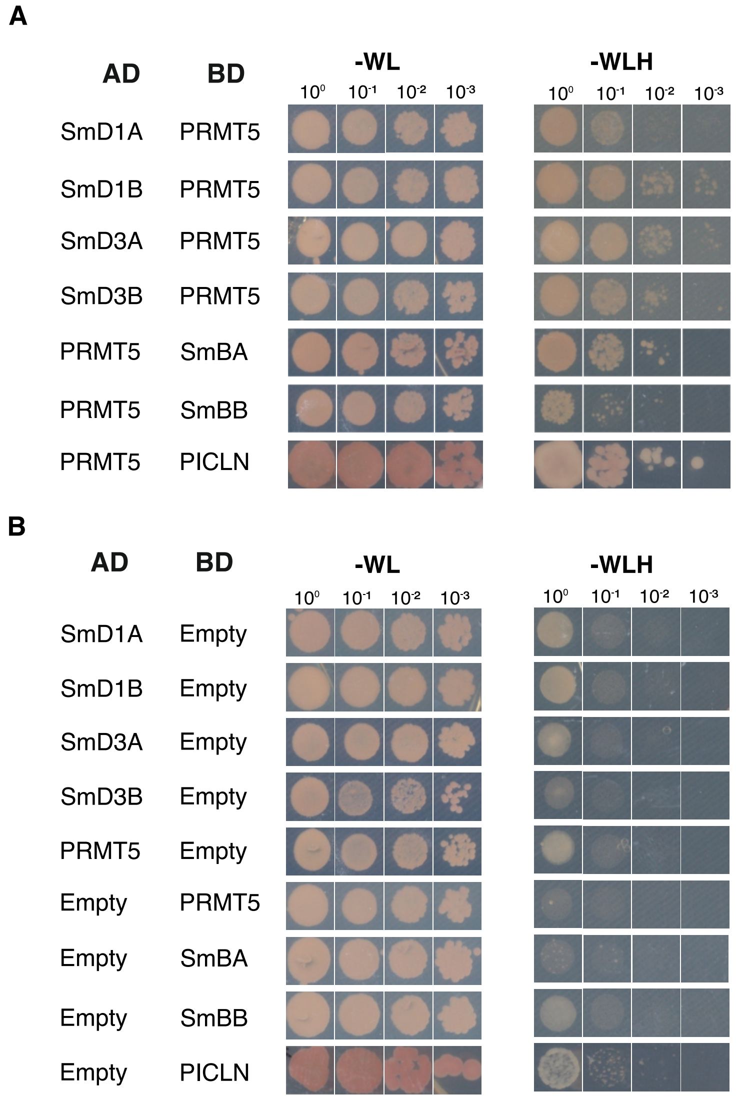
A
AD
BD
-WL
-WLH
10⁰ 10⁻¹ 10⁻² 10⁻³
SmD1A PRMT5
SmD1B PRMT5
SmD3A PRMT5
SmD3B PRMT5
PRMT5 SmBA
PRMT5 SmBB
PRMT5 PICLN
B
AD
BD
-WL
-WLH
SmD1A Empty
SmD1B Empty
SmD3A Empty
SmD3B Empty
PRMT5 Empty
Empty PRMT5
Empty SmBA
Empty SmBB
Empty PICLN

**Figure S10**

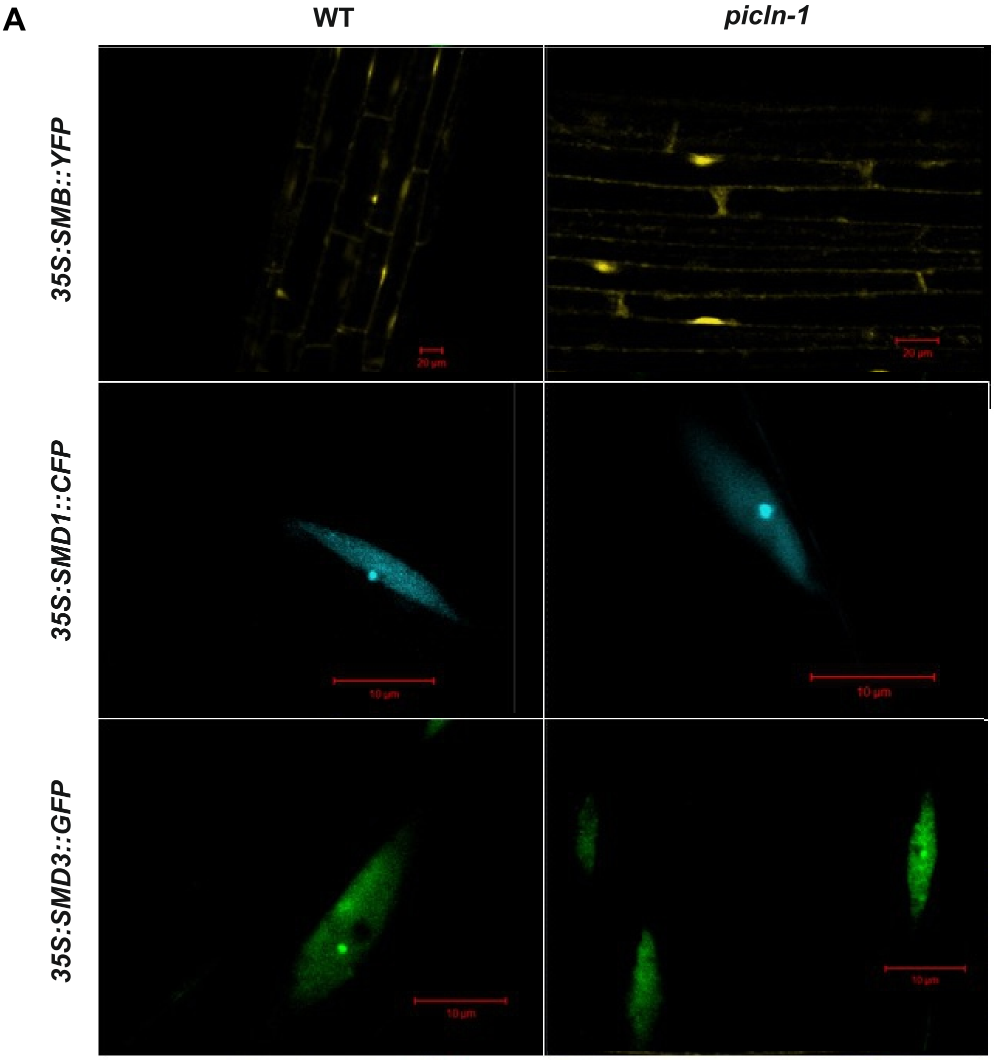
A
WT
picln-1
35S:SMB::YFP
35S:SMD1::CFP
35S:SMD3::GFP
20 µm
10 µm

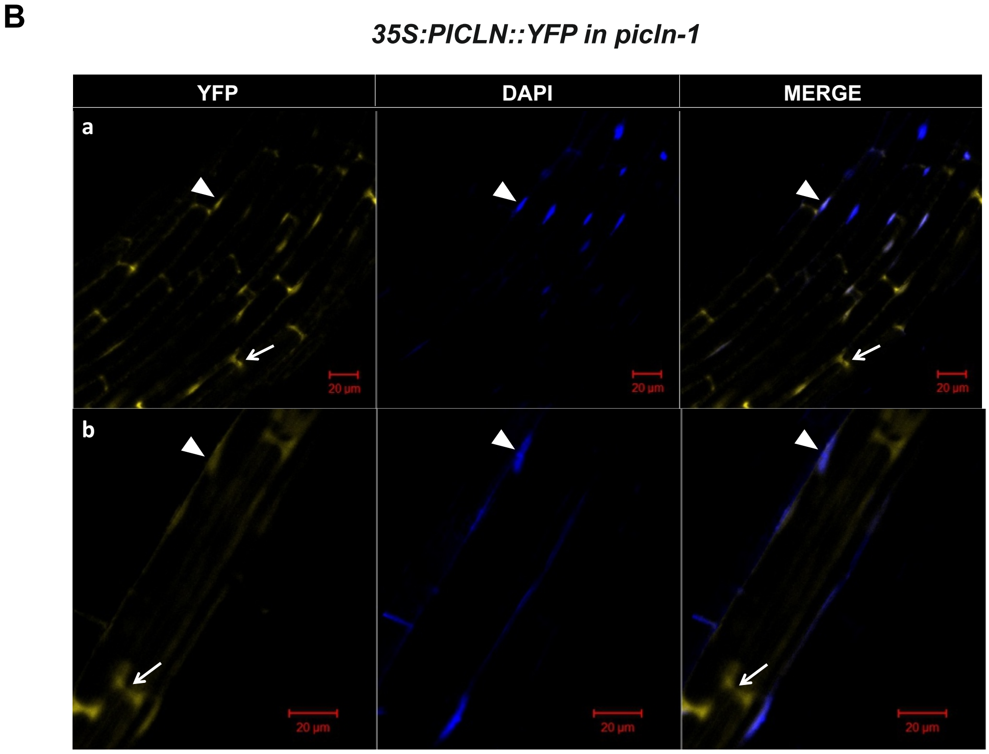
B
35S:PICLN::YFP in picln-1
YFP
DAPI
MERGE
a
b
20 µm

Figure S11

**A**

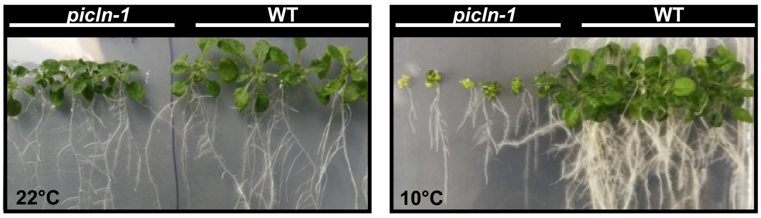


**B**

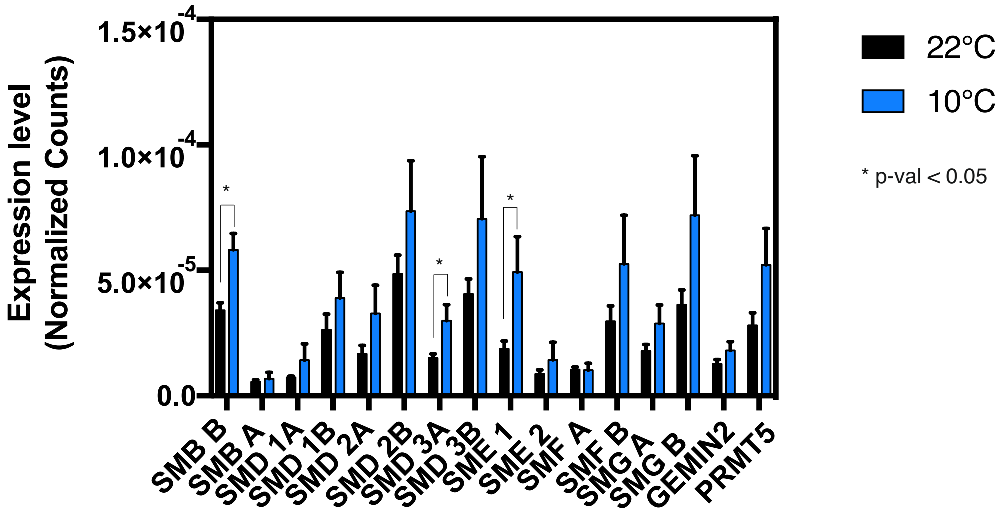


**Figure S12**
